# Identification of arginine-vasopressin receptor 1a (Avpr1a/AVPR1A) as a novel candidate gene for chronic visceral pain

**DOI:** 10.1101/2023.12.19.572390

**Authors:** Leena Kader, Adam Willits, Sebastian Meriano, Julie A. Christianson, Jun-Ho La, Bin Feng, Brittany Knight, Gulum Kosova, Jennifer Deberry, Matthew Coates, Jeffrey Hyams, Kyle Baumbauer, Erin E. Young

## Abstract

Chronic abdominal pain in the absence of ongoing disease is the hallmark of disorders of gut-brain interaction (DGBIs), including irritable bowel syndrome (IBS). While the etiology of DGBIs remains poorly understood, there is evidence that both genetic and environmental factors play a role. In this study, we report the identification and validation of *Avpr1a* as a novel candidate gene for visceral hypersensitivity (VH), a primary peripheral mechanism underlying abdominal pain in DGBI/IBS. Comparing two C57BL/6 (BL/6) substrains (C57BL/6NTac and C57BL/6J) revealed differential susceptibility to the development of chronic VH following intrarectal zymosan (ZYM) instillation, a validated preclinical model for post-inflammatory IBS. Using whole genome sequencing, we identified a SNP differentiating the two strains in the 5’ intergenic region upstream of *Avpr1a*, encoding the protein arginine-vasopressin receptor 1A (AVPR1A). We used behavioral, histological, and molecular approaches to identify distal colon-specific gene expression differences and neuronal hyperresponsiveness covarying with *Avpr1a* genotype and VH susceptibility. While the two BL/6 substrains did not differ across other gastrointestinal (GI) phenotypes (e.g., GI motility), VH-susceptible BL/6NTac mice had higher colonic *Avpr1a* mRNA and protein expression. Moreover, neurons of the enteric nervous system were hyperresponsive to the AVPR1A agonist AVP, suggesting a role for enteric neurons in the pathology underlying VH. These results parallel our findings that patients’ colonic *Avpr1a* mRNA expression was higher in patients with higher pain ratings. Taken together, these findings implicate differential regulation of *Avpr1a* as a novel mechanism of VH-susceptibility as well as a potential therapeutic target specific to VH.

**Summary:** A combination of approaches, from genomic analysis to functional analyses, confirm *Avpr1a* as a high priority candidate gene for visceral pain.

## Introduction

Irritable bowel syndrome (IBS) is one of the most common disorders of gut-brain interaction (DGBIs). Patients report persistent abdominal pain in the absence of structural abnormalities or ongoing inflammation [2–4], along with alterations in bowel habits that can include diarrhea, constipation, or a combination of the two. Prior work suggests that only 30% of people with DGBIs will seek medical care, but patients with unexplained abdominal pain are 40% more likely to receive an opioid prescription than patients with structural GI disorders (e.g., inflammatory bowel disease)[5]. Notably, approximately 60% of patients with an opioid use disorder (OUD) report the presence of at least one chronic pain condition prior to their first OUD diagnosis[6]. This reflects both the lack of visceral pain-specific therapeutic options and the current knowledge gap surrounding the underlying mechanisms of DGBI-related visceral pain.

Visceral hypersensitivity (VH) refers to increased sensitivity to mechanical stimulation of the visceral organs (e.g., bowel) and is a driver of chronic abdominal pain in DGBIs. VH of the distal colon/rectum occurs in up to 90% of patients with IBS[7, 8], so a valid preclinical model should manifest VH in the absence of ongoing inflammation and structural disease of the bowel[9]. In contrast to other preclinical chemical models of VH[10–18], intracolonic instillation of zymosan (ZYM), a protein-carbohydrate complex found in the cell walls of the yeast *Saccharomyces cerevisiae*, results in VH in the absence of structural damage or ongoing inflammation, paralleling the clinical presentation of IBS[1]. ZYM-induced VH has been studied preclinically using both rat and mouse models[17, 18], yet differential susceptibility among substrains has yet to be fully leveraged to explore genetic determinants of VH susceptibility. To identify the contribution of novel genes/genetic variants involved in the susceptibility of VH, we compare the responses of two murine substrains that differ genetically due to the elimination of large portions of the genome; otherwise, biologically, the two strains are similar [19–23].

Genetic studies evaluating IBS have focused primarily on predictors of the diagnosis or associations between polymorphisms and IBS-subtype classifications but have not explicitly evaluated the genetic underpinnings of abdominal pain persistence and/or severity. Given that abdominal pain represents the key symptom across all IBS subtypes[24], understanding the genetic contribution to VH may provide insight into novel therapeutic targets for treating abdominal pain in this population. Capitalizing on differential susceptibility to ZYM-induced VH between two commonly used C57BL/6 (BL/6) substrains (BL/6J from Jackson, BL/6NTac from Taconic) and whole genome sequencing[23], we identified arginine-vasopressin receptor 1A (*Avpr1a*) as a high priority candidate gene for VH. In this study, we use the ZYM-VH model using molecular, electrophysiological, and calcium (Ca^2+^) imaging approaches to identify whether differential expression of *Avpr1a* corresponds to alterations in specific neuronal responses between strains, supporting a role for *Avpr1a*/AVPR1A in VH susceptibility. We additionally measure the relationship between visceral pain intensities experienced and colonic *AVPR1A* expression using colorectal biopsies from patients. Our findings propose *Avpr1a* as a novel visceral pain specific VH therapeutic target and a potential VH susceptibility biomarker in DGBIs.

## Materials and Methods

### Subjects

#### Mice

Adult (8 – 10-week-old) male C57BL/6 mice from Jackson Labs (C57BL/6J; BL/6J) and Taconic (C57BL/6NTac; BL/6NTac) were housed at the University of Pittsburgh and University of Kansas Medical Center in animal facilities on a 12:12 light: dark cycle with food and water available *ad libitum*. Animal use protocols conformed to NIH guidelines and were approved by the Institutional Animal Care and Use Committees at the University of Pittsburgh and the University of Kansas Medical Center and conformed to the Committee for Research and Ethical Issues of IASP. <u>Human subjects.</u> Outpatients presenting for an elective colonoscopy at the University of Pittsburgh Medical Center were approached about participation. To be included in this study, participants had to be between 18 and 70 years of age, able to understand, speak, and write English, and make their own health care decisions. Written informed consent that was HIPAA compliant and approved by the University of Pittsburgh IRB (protocol no 11070169) was obtained for all participants.

### ZYM model of VH

Intracolonic treatment with saline (SAL) or zymosan (ZYM) was performed daily for three consecutive days as described previously[25]. Mice were anesthetized with isoflurane gas anesthesia (2% for induction followed by 0.50-1.0% for maintenance) followed by a transanal administration of 0.1 mL of SAL or ZYM (30 mg/ml, Sigma-Aldrich, St. Louis, MO) solution via a 22-gauge gavage needle. Mice were inverted slightly for 60 seconds following installation to prevent leakage following administration and then returned to the home cage for recovery from anesthesia. Note that for **Figure 1**, ZYM-treated mice were compared to their own Naïve (no SAL) baseline scores prior to treatment with ZYM.

**Figure 1.**
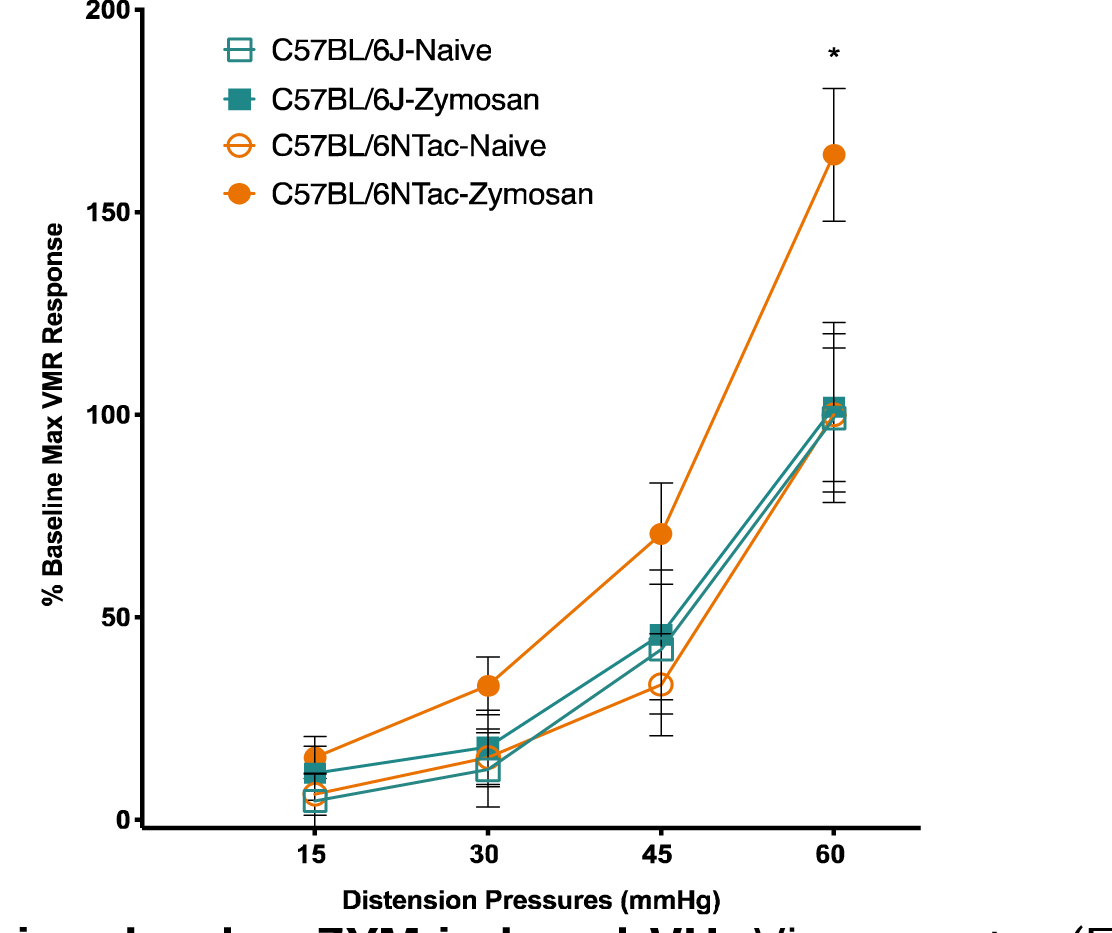
C57BL/6 mice develop ZYM-induced VH. Visceromotor (EMG) responses were measured during colorectal balloon distension pre- and post-intracolonic zymosan (ZYM) instillation for each strain. The graph shows the following groups: C57BL/6J- Naïve (open teal square), C57BL/6J-ZYM (teal square), C57BL/6NTac-Naïve (open orange circle), and C57BL/6NTac (orange circle) mice. Each strain was normalized to its own naïve (baseline) group. Only VMR to 60 mm Hg distention pressure confirmed a significant main effect of condition [F (3, 38)] = 3.493, p < 0.05).

### Visceromotor response to colorectal distension

Evaluation of sensitivity to colorectal distension (CRD) was conducted at baseline (prior to ZYM instillation) and again 21 days after the final ZYM installation based on prior work identifying this as a time when VH is established[17, 26, 27]. Mice (n=7-10/condition) were anesthetized with isoflurane gas and the lower abdomen was shaved and washed with betadine, followed by 70% ethanol. A vertical skin incision, approximately two centimeters in length, was made in the lower abdomen to expose the abdominal musculature, and the skin was separated from the underlying muscle around the side of the abdomen. A pair of stainless-steel wire electrodes were implanted in the right abdominal musculature and then secured with sutures (7-0 Prolene) at the site of contact with the muscle and secured in place by a common suture anchor to the muscle wall of the lower abdomen. The free ends of the electrode wires were tunneled subcutaneously along the side of the abdomen internally and externalized at the nape of the neck, where the ends were anchored with internal sutures for access during testing. Both the abdominal and neck incisions were sutured (4-0 Vicryl), the wounds cleaned, and subjects were administered a subcutaneous injection of 2 mg/kg of buprenorphine in sterile saline. Buprenorphine was administered every 12 hours for the first 48 hours post-surgery (a total of four administrations), and mice were monitored for signs of distress. Mice were placed in the home cage on a heating pad set to low and monitored for one-hour post-surgery to ensure recovery. One week after electrode implantation, mice were transferred to a separate room for visceromotor response testing. Under light anesthesia (inhaled isoflurane, 2%), a custom-made, catheterized polyethylene balloon (1.5 cm long, ∼0.9 centimeters) was inserted through the anus until the proximal end of the balloon was 0.5cm from the anal verge (total balloon insertion, 2cm) and secured to the base of the tail with tape. The mouse was then placed into a plastic cylinder to limit movements, and the free ends of the electrode wires were attached to a differential amplifier (Model 1700, A-M Systems, Sequim, WA). Mice were allowed to recover from anesthesia for 30 minutes. Then electromyographic (EMG) activity, the visceromotor response (VMR), was recorded using Spike 2 software (Cambridge Electronic Design, Cambridge, UK) for 10 seconds before (resting) and during 10 seconds of phasic colorectal distension (15, 30, 45, or 60 mmHg). Mice underwent three trials at each pressure with a four-minute intertrial interval. Responses to distension were quantified as the total area under the curve (AUC) of electromyographic activity during balloon inflation minus resting activity. The three responses to a given distension pressure were averaged for use in all statistical analyses as previously reported[17]. All scores for each mouse were normalized to their maximum response magnitude at baseline, reflecting our focus on differential sensitivity to VH and controlling for baseline differences in VMR to CRD.

### Genomic comparison of pain- and nociception-related genes between C57BL/6J and C57BL/6NTac

A systematic mini review of all genes where variation had previously been associated with pain, hypersensitivity, and/or nociception was performed. This included individual candidate genes and genes from genomic loci identified in the mouse (e.g., QTLs). This gene list was then compared to a list of all differential single nucleotide variants (SNVs) between the BL/6NTac and BL/6J SNPs provided from recently published work by Mortazavi and colleagues[23]. Data and code availability of whole genome sequencing and differential SNP data can be found under the “Key resources table” in the deposited data section[23]. Differential SNPs were then identified to either be within a known gene sequence (i.e., 5’ flanking region, exon, introns, or 3’ UTR) or near a known gene (defined as 18 kb upstream and downstream of the coding sequence) using Ensembl Archive Gene Browser [Nov-2020; Genome assembly GRCm38.p6 (GCA_000001635.8)]. We then generated a finalized candidate gene list where variation(s) within pain/nociception genes differentiated the two strains. Refer to **Figure 2** for a diagram of our review workflow.

**Figure 2.**
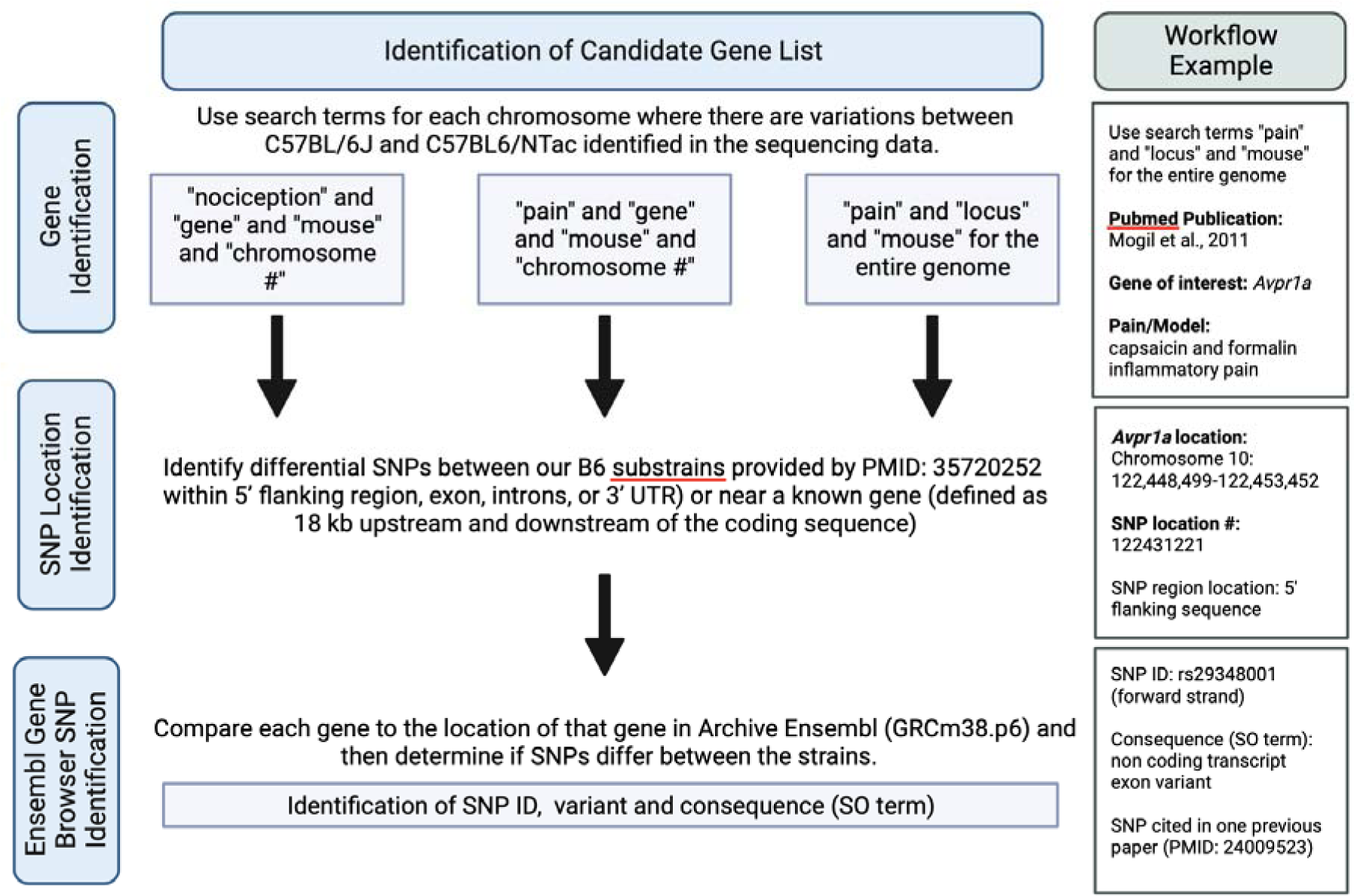
Workflow for identifying pain candidate genes from whole genome sequencing data. In brief, we performed a systematic review and collected a list of genes related to visceral pain, nociception, and/or pain assays. From this list, we used whole genome sequencing data published in Mortazavi, Ren [23] to compare the location of our genes of interest to locations of possible differential SNPs between BL/6J and BL/6NTac. It is important to note Archive Ensembl was used to match and compare whole genome sequencing data to the appropriate gene loci. To the right of the general flow is an example workflow used to identify an intergenic SNP within the 5’ region of *Avpr1a*.

### Histology and Immunohistochemistry

Mice (n=5/condition) were overdosed with inhaled isoflurane (> 5%) and perfused transcardially with ice-cold Hanks Balanced Salt Solution (HBSS) on day 21 post-SAL or ZYM treatment. Approximately 4.5 cm of the distal colon was collected and post-fixed in 10% neutralized formalin fixative. After fixation, the distal colon tissue was embedded in paraffin wax, cut in 8μm thick cross-sections using a microtome, and mounted on glass slides. Sections were deparaffinized, rehydrated, and washed briefly in distilled water. For Hematoxylin and Eosin staining, slides were stained in Harris hematoxylin solution for eight minutes and subsequently washed in running tap water for five minutes. Samples were differentiated in 1% acid alcohol for 30 seconds and washed in running tap water for one minute. Samples were then placed in 0.2% ammonia water or saturated lithium carbonate solution for 30 seconds to one minute and then washed in running tap water for five minutes. Afterward, slides were rinsed in 95% alcohol for approximately ten dips and then counterstained in eosin-phloxine solution for thirty seconds to one minute. Lastly, slides were dehydrated and mounted with a xylene-based mounting medium. For Alcian Blue Staining, slides were stained for goblet cells and mucins using an Alcian Blue dye and Nuclear Red Fast counterstain kit (Abcam; ab150662). Quantification of the percent area of Alcian blue staining, which calculated the percent area of the Alcian Blue dye compared to the rest of the tissue section, was calculated using the Alcian Blue color deconvolution plugin in ImageJ (ImageJ 1.53k Java 1.8.0_172).

### Immunofluorescent staining (PGP9.5 & AVPR1A)

On day 21 post-SAL or ZYM treatment, adult male mice (n=5/condition) were overdosed with inhaled isoflurane followed by intracardiac perfusion of ice-cold HBSS. About 4.5 cm of the distal colon was collected and post-fixed in 10% neutralized formalin fixative. After fixation, the distal colon tissue was embedded in paraffin wax, cut in 8μm thick cross-sections using a microtome, and mounted on glass slides. After sections were deparaffinized, rehydrated, and washed briefly in distilled water. Formalin-fixed tissue sections were deparaffinized and rehydrated. Blocking solution (5% donkey serum in PBST) was applied for twenty minutes at room temperature. Primary antibodies were applied in blocking solution overnight in a humidified, light-protected chamber at 4°C (1:1500 anti- PGP9.5 & 1:1000 anti-AVPR1A). The next day, slides were washed with PBST for 15 minutes each (3X). A secondary antibody cocktail was then applied (donkey anti-rabbit 555-red - PGP9.5 and Donkey anti-mouse 488-green – AVPR1A in PBST) for one hour in a light-protected chamber at room temperature. Tissue sections were then washed for 15 minutes each time (3X). DAPI and Vectashield mounting media were applied and covered with a coverslip. Images were taken on an 80i fluorescent scope using the DAPI, GFP, and Cy3 filter settings. Images were merged and processed using ImageJ[28].

### Gastrointestinal transit (GIT) and colonic transit (CT)

Experimental measures for colonic motility were based on previous experimental design[29]. Mice (n=5/condition) were lightly anesthetized with 2% isoflurane in oxygen. The mouse’s head was carefully hyperextended to pull out the tongue with blunt forceps. 100 μl fluorescein-labeled dextran (FITC-dextran, 70,000 MW, Sigma Aldrich) was briefly injected with a 1.0 mL syringe oral gavage. While remaining under inhaled 2% isoflurane anesthesia, colonic transit experiments were concurrently performed. A 2 mm glass ball was inserted 3 cm past the anus into the colon using a fistula probe (metal rod, 2 mm diameter). Immediately after removing the fistula probe, mice were placed in a transparent cage, and the time from when the fistula probe was pulled out of the colon to when the glass ball was excreted out was measured. For the remaining 90-minute period, mice were given access to food and water. After 90 minutes, mice were euthanized, and the entire gastrointestinal tract from stomach to rectum. The entire tract was placed on a polystyrene pad to avoid excess stretching.

The total length of the small bowel and the entire length of the colon were measured. The gastrointestinal tract was then transected into 15 prepared segments: the stomach (#1), ten equally sized small bowel segments (#2-11), cecum (#12), and three equally sized colonic segments (#13-15). Each section was luminally flushed with 1 mL of Krebs Henseleit Buffer (KHB) solution, collected in 2 mL tubes. The tubes were centrifuged at 12,000 rcf for five minutes at room temperature. The clear supernatants were transferred to new 1.5 mL tubes and stored in the dark at 4°C until analysis (can be stored for several days at 4°C without significant loss of FITC signal. For more extended storage, include 0.09% Natrium-azide or store at < -18°C). For analysis, 100 μl of each supernatant was pipetted in duplicate onto a black 96-well plate (Greiner Bio One, Frickenhausen, Germany). Duplicate 100 μl KHB served as blanks. The fluorescence/well was quantified at 494 nm (absorption) / 521 nm (emission) wavelength. For both gastrointestinal and colonic transit, the geometric center (GC) of FITC-dextran distribution, the center of gravity for the distribution of the marker, was calculated by the following formula[29]:

GC = ∑ (% of total fluorescent signal per segment * segment number) / 100

### Stool consistency assay

Stool consistency can be used as a quantitative assessment of water content in fecal samples, allowing for objective determination of the presence of altered bowel habits (i.e., diarrhea and/or constipation). Four fecal pellets were collected (immediately after excretion) from each mouse (n=5/condition) and weighed to four decimal places. The pellets were then microwaved for six minutes. Following dehydration, the pellets were weighed again. The wet weight and dry weight were normalized by dividing each weight by the number of pellets. The difference between the normalized wet and dry weights was used in statistical analyses.

### Retrograde labeling of colon-specific extrinsic afferents

Fourteen days after treatment with ZYM (or SAL), mice (n=5/condition) were anesthetized by inhaled isoflurane (4% for induction, 2% for maintenance), and a laparotomy was performed to expose the pelvic viscera. Using a Hamilton microsyringe with a 22-gauge needle, a 15 μl injection of Alexa Fluor 488-conjugated cholera toxin-β (CTB) (2 mg/ml) was made beneath the serosal layer in the distal region of the colon and the abdominal incision sutured. Mice were returned to their home cage for seven days before further experiments.

### Avpr1a mRNA expression in mouse colon and colon-specific extrinsic primary afferents

Mice of both substrains (n=5/condition) were treated with either intrarectal SAL or ZYM for three consecutive days, as described above. *Retrograde labeling of colon-specific afferents.* Fourteen days after the final ZYM instillation, mice were anesthetized with gaseous isoflurane, and a laparotomy was conducted to open the abdominal cavity. Each mouse received a series of injections of Alexa Fluor 488- conjugated cholera toxin B (CTB) (Invitrogen; C-34775). These injections were into the subserosa of the distal colon, and the cumulation of the injections’ total volume was 10 μl per mouse. *Tissue collection.* Seven days later, mice were overdosed with isoflurane and perfused with ice-cold Ca^2+^/Mg^2+^-free Hank’s Balanced Salt Solution (HBSS, Invitrogen). A two-centimeter segment of the distal colon was collected and flash-frozen on dry ice. At the same time, thoracolumbar (T12-L2) and lumbosacral (L5-S1) dorsal root ganglia were rapidly dissected and prepared for culture as described previously[30]. *Cellular Preparation.* Dissociated cells were resuspended in F12 Nutrition Mixture (Invitrogen; REF# 11765-054) containing 10% fetal calf serum and antibiotics (penicillin/streptomycin, 50 U/ml) and plated onto laminin (0.1mg/ml) and poly-D-lysine (5mg/ml) coated glass coverslips. No additional growth factors were added to the culture medium. Cells were incubated overnight at 37°C, and individual CTB-positive colon-specific neurons were collected with large-bore (∼50 μm) glass pipettes and expelled into microcentrifuge tubes containing reverse-transcriptase (RT) mix (Invitrogen). For each experiment, negative controls consisted of omitting reverse transcriptase or using a cell-free bath aspirate as template. The first-strand cDNA from a colonic sensory neuron was used as template in a PCR reaction containing 1×*GoTaq* reaction buffer (Promega Madison, WI), 20 mM outer primers, 0.2 mM dNTPs, and 0.2 mL *GoTaq* DNA polymerase (Promega). Each initial PCR product served as template in a subsequent PCR reaction using a nested primer pair, the products of which were electrophoresed on 2% agarose–ethidium bromide gels and photographed. Only neurons producing detectable amplification of a housekeeping gene (*Gapdh*) were analyzed further. The following primers were used for first-round amplification: *Avpr1a*, 5′-CCTACATGCTGGTGGTGATG-3′ (forward) and 5′-TCTTCACTGTGCGGATCTTG-3′(reverse); *Gapdh*, 5′-AACTTTGGCATTGTGGAAGG-3′ (forward) and 5′- CCCTGTTGCTGTAGCCGTAT-3′ (reverse). For qRT-PCR, the first-strand cDNA of cells expressing the target genes was pre-amplified (26 cycles) using the PCR condition as described above. For second round (nested) amplification, the forward and reverse primer sequences were as follows: *Avpr1a*, 5′-GTCCGAGGGAAGACAGCATC-3′ and 5′-GATCTTGGCACGGGAAATGC-3′, respectively; *Gapdh*, 5′- ATGAATACGGCTACAGCAACAGG-3′ and 5′-CTCTTGCTCAGTGTCCTTGCTG-3′, respectively. Final amplification products had a predicted size of 338 bp (*Avpr1a*) and 301 bp (*Gapdh*) and were separated by electrophoresis. The products were used as a template for quantitative real-time PCR (qRT-PCR) using ABsolute^™^ QPCR SYBR^®^ Green ROX mix (ABgene, Rochester, NY) in an Applied Biosystems (Foster City, CA) 5700 real-time thermal cycler. Colorectal samples only underwent PCR with the second strand (nested) primer set as the pre-amplification of the target product was not necessary for adequate quantification. Threshold cycle (C_t_) values were recorded to measure initial template concentration, and relative fold changes in mRNA levels were calculated using the 2^-ΔΔCt^ method using *Gapdh* as a reference standard.

### Colorectal AVPR1A protein expression

Total protein was isolated from approximately 50 mg of snap-frozen distal-colon tissue using Cell Extraction Buffer (Invitrogen) containing Halt protease and phosphatase inhibitors (ThermoFisher Scientific, Waltham MA) and Na_3_VO_4_. Protein concentrations were determined using the Nanodrop spectrophotometer. Protein levels of AVPR1A were measured using ELISA per the manufacturer’s instruction (MSB, Cat # MSB017824).

### Sensory neuron response to colonic stretch (single fiber recordings, aka fiber teasing)

Detailed procedures were described previously[31]. Briefly, mice (n=5/condition) were killed via CO_2_ inhalation followed by exsanguination after perforating the right atrium. The distal colorectum and pelvic nerves (PN) were dissected and transferred to ice-cold Krebs solution bubbled with carbogen (95% O_2_, 5% CO_2_). The colorectum was opened longitudinally, pinned flat mucosal side up in a tissue chamber, and the PNs extended into an adjacent recording chamber filled with paraffin oil. The tissue chamber was superfused with a modified Krebs solution containing the following (in mM): 117.9 NaCl, 4.7 KCl, 25 NaHCO_3_, 1.3 NaH_2_PO_4_, 1.2 MgSO_4_, 7 H_2_O, 2.5 CaCl_2_, 11.1 D-glucose, 2 butyrate, and 20 acetate at a temperature of approximately 3°C to which was added nifedipine (4 μM) and indomethacin (3 μM). The PN was teased into fine bundles (∼10 μM thickness) for single-fiber recording. A custom-made rake with fine pins (1 mm interval) were inserted along the antimesenteric edge of the colorectum to permit homogeneous, circumferential stretch by a slow ramped force (0—170 mN at 5 mN/s) corresponding to intraluminal pressures of 0 – 45 mmHg. To independently assess the response to longitudinal stretch, separate colorectum samples were rotated 90° prior to insertion of the custom-made claws. After establishing a baseline (control) of the stimulus response function (SRF), the receptive field of the afferent in the colorectum was isolated by a brass ring, and the Krebs solution within was removed and replaced by 150 μl of Krebs solution containing arginine-vasopressin (0.1 μM; Sigma-Aldrich V9879) (five minutes). The tubing was then removed, re-exposing the ending to Krebs solution, and a response to the same stretch (circumferential or longitudinal) was acquired immediately after AVP. Twenty minutes after exposure to the last concentration tested, an additional SRF was recorded to conclude the protocol. Electrical signals generated by PN afferent endings were filtered (0.3 to 10 kHz) and amplified (x10,000) by a low-noise AC differential amplifier (DAM 80; World Precision Instruments, New Haven, CT). The signal was digitized at 20 kHz using a 1401 interface (Cambridge Electronic Design, Cambridge, UK) and stored on a PC. The signal was also monitored online by an audio monitor (Grass AM8; Astro-med, West Warwick, RI). Action potentials were analyzed offline using the Spike 2 wave mark function and discriminated as single units based on principal component analysis of individual spike waveforms. No more than two easily discernible active units in any record were studied to avoid errors in discrimination.

### Clinical subject recruitment, phenotyping, and sample collection

The methods for sample collection and recruitment of ulcerative colitis (UC) patients have been described in detail elsewhere[32]. Individuals with an established diagnosis of UC and healthy controls undergoing colonoscopy at a large, metropolitan, academic medical center were enrolled. All participants rated abdominal pain using a visual analog scale (VAS) and completed standardized surveys addressing anxiety or depression (HADS) and gastrointestinal symptoms (Rome III questionnaire). Patient age, sex, and severity of inflammation (based on endoscopic and histologic assessment) were determined. A subset of study participants used in a previously published study[32] were divided into the following cohorts for analysis of *AVPR1A* mRNA expression (based upon underlying diagnosis and pain status): 1) Healthy Controls, 2) active UC with (abdominal) pain, 3) active UC without (abdominal) pain. In addition to UC patient recruitment and sample collection, we analyzed a subset of pediatric IBS patients undergoing colonoscopy at Connecticut Children’s Medical Center. Pediatric patients (males and females, age 7- 17) undergoing diagnostic colorectal biopsy for abdominal pain were compared (n=3/patient) to pediatric patients undergoing painless non-cancerous polyp monitoring[33]. Patients meeting Rome III criteria for IBS (and controls) in the absence of other diagnoses (e.g., *H. pylori* and IBD) or clinical signs of disease were administered the Pain Burden Inventory (PBI) to measure self-reported pain severity. We evaluated relative *AVPR1A* mRNA expression (2^-ΔΔCt^ method) in biopsies from all patients and compared IBS patients with low and high self-reported pain [defined as a PBI score of 0-2 (low pain, n=13) or greater than a PBI score of 2 (high pain, n=25), respectively].

### Assessment of AVPR1A mRNA expression in human rectal biopsies

To assess the translational potential of this relationship between *Avpr1a* expression and persistent abdominal pain, we evaluated *AVPR1A* mRNA expression in colorectal biopsies previously collected from two cohorts of patients with UC and IBS and comparing those with pain to those reporting no pain at the time of diagnosis[32]. If *AVPR1A* expression contributes to persistent visceral VH/abdominal pain in humans, as in our mouse model, then expression should differ between patients with and without pain. Rectal biopsies were obtained and processed using qRT-PCR to determine the transcript expression of *AVPR1A* (PrimerBank ID: 33149325c1). Total RNA was extracted using RNeasy isolation columns (Qiagen, Hilden, Germany) and real-time quantitative PCR was undertaken as described previously[34]. According to manufacturer’s instructions, mRNA samples underwent reverse transcription with iScript cDNA synthesis kits (BioRad, Hercules, CA). Quantification with a nana-drop and loading of 2 ng of total cDNA were placed into each reaction. cDNA template was then used for qRT-PCR SSO Universal SYBR^®^ Green Master Mix with ROX (BioRad, Hercules, CA) in an Applied Biosystems StepOne Plus PCR machine (Foster City, CA). Threshold cycle (C_t_) values were recorded as a measure of initial template concentration, and relative fold changes in mRNA levels were calculated by the ΔΔC_t_ method using GAPDH as a reference standard: GAPDH, 5’-AAG-GAC-TCA-TGA-CCA-CAG-TTC-ATG-3’ (forward strand) and 5’-TTG-ATG-GTA-CAT-GAC-AAG-GTG-CGC-3’ (reverse strand). Primer sets were designed using BLAST. All primer sets were evaluated for efficiency and were found to have > 95% amplification of the template sequence at 1 uM concentration. Single sequence amplification was confirmed by running melt curve analysis on all PCR products, and all products underwent gel electrophoresis to confirm a single amplified product of the expected size. The following primers were used for amplification: AVPR1A, 5’-TGT-AAA-ACG-ACG-GCC-AGT-CCC-CAG-AGT-TAA-GAC-AGT-TGC-3’ (forward strand), and 5’-CAG-GAA-ACA-GCT-ATG-ACC-CCG-GTT-TAC-CCT-TGC-ACT-TT-3’ (reverse strand).

### Dissociation, culture, and imaging

Seven days after injection of retrograde tracer, mice were overdosed with inhaled isoflurane (>5%) and transcardially perfused with 4°C Ca^2+^/Mg^2+^-free 1X HBSS (Invitrogen). Bilateral T11-L2 DRGs, L3–S1 DRGs, and nodose (ND) ganglia were dissected and placed separately into 1X HBSS for cellular dissociation. The enteric nervous system was collected from a separate cohort of mice. The myenteric plexus was collected and dissociated as described elsewhere[35, 36]. Briefly, 1-2 cm of the distal colon was collected per mouse and placed in ice-cold 1X HBSS. Colon samples were mounted to an agar plate surface while submerged in 1XHBSS. Additional mesenteries, fat tissue, and fecal matter were removed. The mucosa layer, followed by submucosal layer were carefully removed to easily isolate enteric neurons from the myenteric plexus. Isolated myenteric tissue was placed in culture tubes for cellular dissociation. Enzymatic dissociation of smooth muscle and myenteric plexus tissue was performed as described in previous work[35, 36]. After cellular dissociation of each tissue sample (DRG, ND, Enteric) (as stated above), cells were resuspended in F12 media (Gibco) containing penicillin/streptomycin (50U/ml) and 10% FBS and plated onto 12mm Poly-D-Lysine/Laminin pre-coated coverslips (BD Biosciences, Franklin Lakes, NJ). Cells were incubated overnight at 37°C and imaged the following day (sixteen to twenty-four hours after plating cells) as published elsewhere[37]. Before imaging, each coverslip was incubated in 3 μl fura-2 acetoxymethyl (AM) ester (Invitrogen) in 1X HBSS containing 5 mg/ml BSA (Sigma-Aldrich) for 30 minutes at 37°C. Coverslips were placed in a QE-1 quick change platform (Warner Instruments, Hamden, CT) and mounted on a Nikon Eclipse Ti inverted microscope stage with a constant flow of 1X HBSS buffer that was controlled by a gravity flow system (Warner Instruments). Perfusate temperature was maintained at 30°C, and all agonists were dispensed with a gravity-feed pinch valve control perfusion system and included 30 mM K^+^ (High K^+^), capsaicin (Sigma-Aldrich, 1uM), and Arginine-Vasopressin (AVP, Millipore Sigma V9879, 0.1 uM). Capsaicin was dissolved in 1-methyl-2-pyrrolidinone to create a 10 mm stock solution; 1.0 μM capsaicin was made fresh daily in 1X HBSS. Firmly attached, CTB-positive neurons were identified as regions of interest in the software (Nis-Elements Version 4.60). Unlabeled, adjacent cells were also identified and imaged. Absorbance data at 340nm and 380nm were collected at 1Hz during drug application using a pco.Edge 4.2 LT camera (PCO-TECH, Romulus, MI). Responses were measured as the ratio of 340/380nm excitation and 510nm emission controlled by a high-speed Fura/widefield Xenon illuminated filter wheel (Boyce Scientific). All fields were first tested with a brief application (5s) of 30 mm K^+^ (high K^+^) and only cells responsive to High K+ were identified as healthy and responsive and included in subsequent analyses. Five minutes after application of high K^+^, capsaicin was applied until cells started to show peak fluorescence responses (typically ∼2 seconds after the application start), followed by a 10-minute washout period prior to AVP application. AVP is applied via a micropipette in droplet intervals until cells started to show peak fluorescence responses (typically shown immediately after a couple of droplets are placed onto the coverslip). Once an increased change in fluorescence occurs, the agonist application of each agonist is stopped, and a wash buffer is turned on for ∼5 seconds. Peak Ca^2+^ influx was calculated using MATLAB, and responses > 0.1 ΔF_340/380_ were included in the analysis. The prevalence of capsaicin- or AVP-responsive neurons was determined as a percentage of total healthy (High K^+^-responsive) CTB-positive and unlabeled cells for both DRG and ND ganglia. All responsive enteric neurons were included in the analyses.

### Statistical analysis

Outcomes from VMR were analyzed using a 2 (BL/6NTac vs BL/6J) x 2 (pre-ZYM vs. post-ZYM-VH) repeated measure comparison. Post-hoc group comparisons using Fisher’s LSD confirmed that the VMR to 60 mm Hg distention pressure differed between B/6NTac-ZYM and all other conditions RT-PCR, ELISA, single fiber recordings, and Ca^2+^ imaging were analyzed using 2 (BL/6NTac vs BL/6J) x 2 (ZYM vs SAL) analysis of variance (ANOVA) followed by Bonferroni t-tests for critical post-hoc comparisons. This is followed by both a Levene Test for Equality of Equal Variance and a Bonferroni post-hoc for further statistical comparison. Histological stains were evaluated for the percent of colocalization with markers of other cell types (neuronal, muscle, etc.) to determine the distribution of AVPR1A within the heterogeneous cell makeup of the colon wall using programs such as ImageJ/ FIJI. For both clinical cohorts, fold differences in expression compared to the healthy control condition were analyzed using One Way ANOVA (p < 0.05). All data are presented as mean +/- SEM.

## Results

### C57BL/6 substrains exhibit differential susceptibility to VH following ZYM administration

Visceral sensitivity was first determined for BL/6J and BL/6NTac mice by measuring the VMR during colorectal distention before ZYM treatment and 21 days after the final ZYM treatment in the same cohort of mice. A 2 (BL/6NTac vs. BL/6J) x 2 (Pre- vs. Post- ZYM exposure) Analysis of Variance (ANOVA) on the cumulative AUC across all distension pressures revealed a significant main effect of strain (F (1, 11), p = 0.015) and a significant strain x time interaction (F (1, 11) = 6.863, p = 0.024). This effect appeared to be primarily driven by hypersensitivity to CRD at the highest distension pressure (see **Figure 1**). Analysis using a One-Way ANOVA considering only VMR to 60 mm Hg distention pressure confirmed a significant main effect of condition [BL/6J-Naive (Baseline), BL/6J-ZYM, BL/6NTac-Naïve (Baseline), BL/6NTac-ZYM] (p < 0.05). Post-hoc group comparisons using Fisher’s LSD also confirmed that the VMR to 60 mm Hg distention pressure differed between B/6NTac-ZYM and all other conditions (all p < 0.05). No other significant differences were present.

### Weight change, gastrointestinal transit (GIT) and colonic transit (CT) do not differ between treatment and/or substrain

No significant differences were found for colonic- (**Supplemental Figure 2**) or gastrointestinal transit (**Supplemental Figure 3**) assays between the two substrains (BL/6J and BL/6NTac) or treatment condition (ZYM and SAL) (all p > 0.05). There were also no significant differences in fecal water retention between BL/6J and BL/6NTac mice regardless of condition (all p > 0.05) (**Supplemental Figure 4**). Though not statistically significant, the possibility of ZYM causing a low-level diarrhea could be clinically relevant, so we measured body weight to assess dehydration and calorie absorption which could be affected by diarrhea. We found no significant difference in body weight over the development of VH (all p > 0.05) **Supplemental Figure 5**).

### Distal colorectal morphology (inflammation, Goblet cell and mucus production) does not differ between strains

Hematoxylin and eosin (H&E) staining reveals no gross differences between the strains or conditions at Day 21 post ZYM installation, like previous reports that the ZYM inflammation resolves quickly[38]. Representative figures are seen in **Supplemental Figure 6**. Alcian Blue staining, which visualizes acidic epithelial and connective tissue mucins, allowing the measurement of the percent surface area of mucin cell bodies [39], reveals no changes in percent mucin area regardless of strain or condition (all p > 0.05) (**Supplemental Figure 7**).

### Positional identification of Avpr1a as a VH candidate gene

Various phenotypic differences have been reported among C57BL/6 substrains with findings related to learning and memory performance, glucose metabolism, and drug/alcohol responses[40, 41]. Prior work using low-resolution SNP genotyping[42] identified 12 (out of 1449) single nucleotide polymorphisms (SNPs) that differed between BL/6J and BL/6NTac mice[21]. Of these 12 SNPs, three were located on distal chromosome 10, a region previously identified in two separate quantitative trait loci (QTL) for inflammatory pain sensitivity using an injection of formalin into the hind paw[43] and peritoneal acetic acid administration[44]. These two QTLs converge on a single candidate gene from Chromosome 10, arginine-vasopressin receptor 1a (*Avpr1a*), which has subsequently been validated/confirmed as contributing to differences in somatic pain sensitivity in both human and animal studies[45]. Extending this promising finding[42], we leveraged recently published whole genome sequencing of single nucleotide variants (SNV) between BL/6 substrains[23, 46] and identified 5 genes where SNPs differentiated the VH-susceptible and VH-resistant BL/6 substrains (refer to **Figure 2** for a diagram of our gene database workflow). Of these 5 candidates, only *Avpr1a* was associated with variability in more than one pain assay or measure (**Table 1**). We identified a SNV in the 5’ flanking sequence for *Avpr1a*, a region heavily implicated in regulating *Avpr1a* expression[43, 44]. Moreover, we found that this SNV represents an exon variant within an uncharacterized long intergenic non-coding RNA (lincRNA), further implicating a regulatory role for this variant. As such, *Avpr1a* was identified as our highest-priority candidate gene for VH.

**Table 1.**
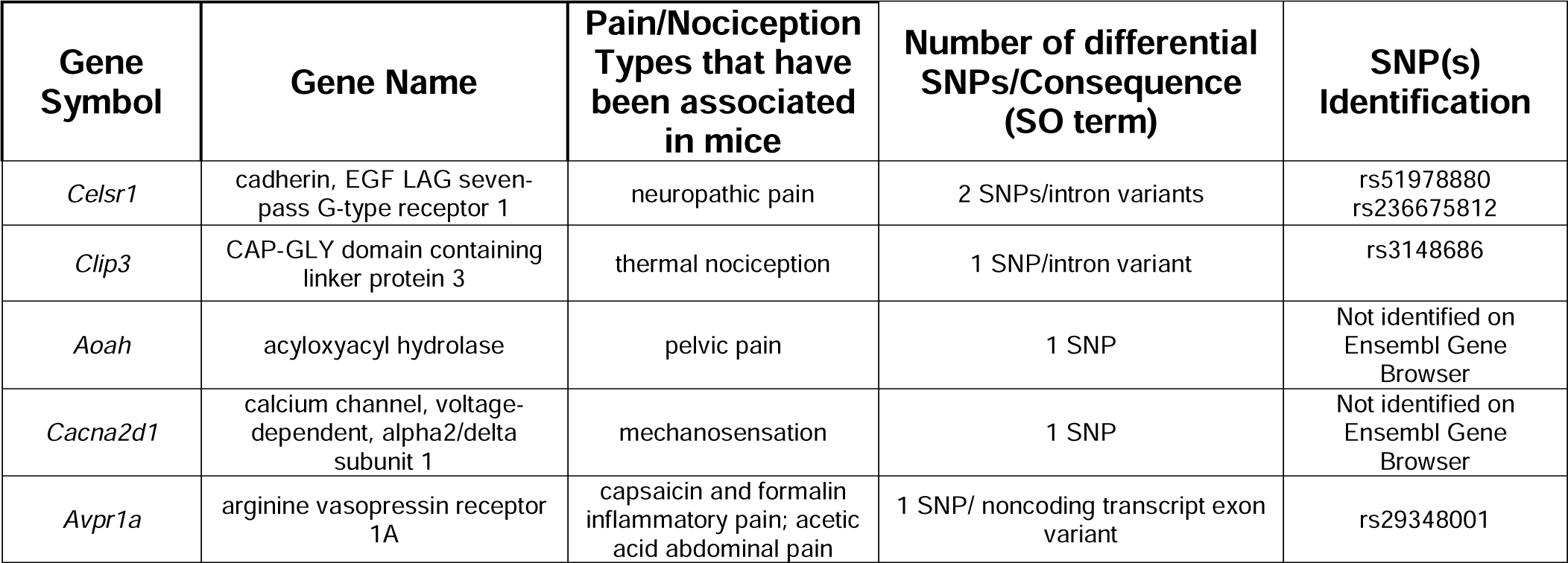
Prioritized candidate genes were identified by comparing BL/6NTac and BL/6J substrains. Only genes where a SNP exists within the 5’ flanking region, exon, introns, or 3’ UTR were included, which were defined as the gene and the surrounding DNA sequence up to 18K nucleotides outside the coding sequence on Ensembl Gene Browser. Note that only *Avpr1a* has been associated with more than one type of pain.

### Increased Avpr1a expression in colorectum but not colon-specific extrinsic primary sensory neurons correspond to VH

*Avpr1a* mRNA expression was determined in both homogenized colorectum and colon-specific primary sensory neurons from BL/6J and BL/6NTac mice, 21 days after SAL or ZYM treatment. *Avpr1a* expression in the colorectum (**Figure 3A**), but not in colon - DRG neurons was increased in BL/6NTac mice treated with ZYM compared to all other conditions. No significant main effects or interactions on the expression of *Avpr1a* in either thoracolumbar or lumbosacral colon-specific primary sensory afferents were detected in a similar 2 x 2 ANOVA (all F [(1, 14)] < 0.831, all p > 0.05) (**Supplemental Figure 1**). In agreement with increased *Avpr1a* mRNA expression in colorectum, AVPR1A protein levels were significantly higher in BL/6NTac mice compared to BL/6J mice 21 days after ZYM (p < 0.05; **Figure 3B**).

**Figure 3.**
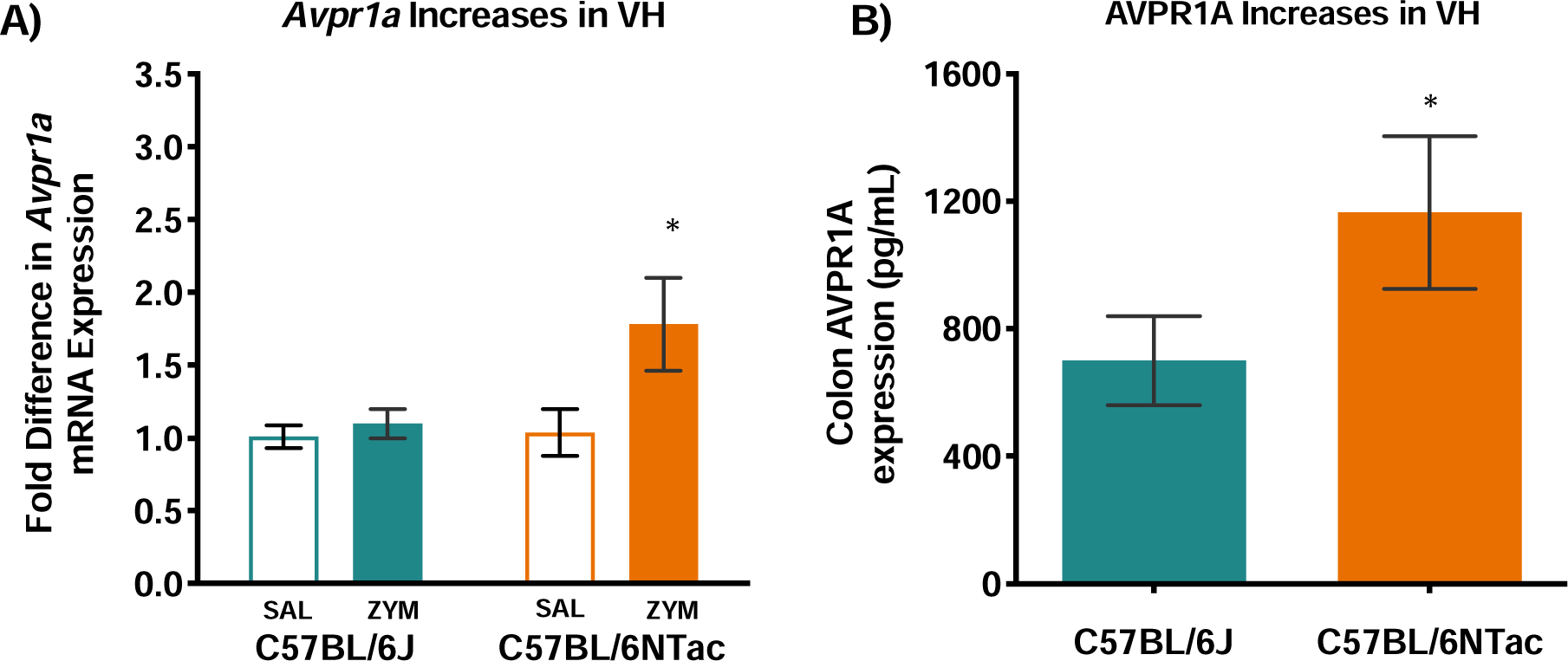
**(A**) **Colonic *Avpr1a* mRNA levels were measured in SAL or ZYM-treated BL/6J and BL/6NTac mice.** A 2 (strain) x 2 (treatment) ANOVA confirmed a significant Strain x Treatment interaction (F (1, 10) = 5.851, p = 0.036). No other significant main effects or interactions were observed (all F < 2.097, all p> 0.05). (**B)** AVPR1A protein levels were measured in ZYM-treated BL/6NTac and BL/6J mice. No significant main effects or interactions on the expression of *Avpr1a* in either thoracolumbar or lumbosacral colon-specific primary sensory afferents were detected in a similar 2 x 2 ANOVA (all F [(1, 14)] < 0.831, all p > 0.05).

### Functional sensitivity to the AVPR1A agonist AVP (arginine-vasopressin) differs between C57BL/6 substrains following ZYM administration

Single fiber recordings revealed a significant increase in sensory neuron activity to longitudinal and circumferential colonic stretch in BL/6NTac mice treated with ZYM when the AVPR1A receptor agonist arginine-vasopressin (AVP) was added to the bath solution (**Figure 4**). Both AUC for the entire ramped stretch stimulus and the sensory neuronal response magnitude to peak stretch force (170mN) was recorded before and after the application of AVP. Specifically, BL/6NTac-ZYM mice had greater AUC compared to BL/6J-ZYM mice and longitudinal stretch in BL/6NTac-ZYM mice elicited the greatest response magnitude (highest AUC) compared to all other groups (p < 0.05). (**Figure 4A**). Similarly, 2 x 2 ANOVA for response during peak stretch force application confirmed a significant main effect of Strain confirming a greater response magnitude in BL/6NTac mice treated with ZYM compared to BL/6J mice treated with ZYM (**Figure 4B**). No other significant main effects or interactions were observed (all p < 0.05). Still, these data align with the strain difference in susceptibility to the VH phenotype even though the maximum stretch force used here corresponds to lower colorectal distension pressure than those administered when measuring VH by VMR to CRD. These data confirm that the increased *Avpr1a*/AVPR1A expression has functional consequences for the response to colorectal stretch corresponding to increased expression and behavioral hypersensitivity in BL/6NTac mice.

**Figure 4.**
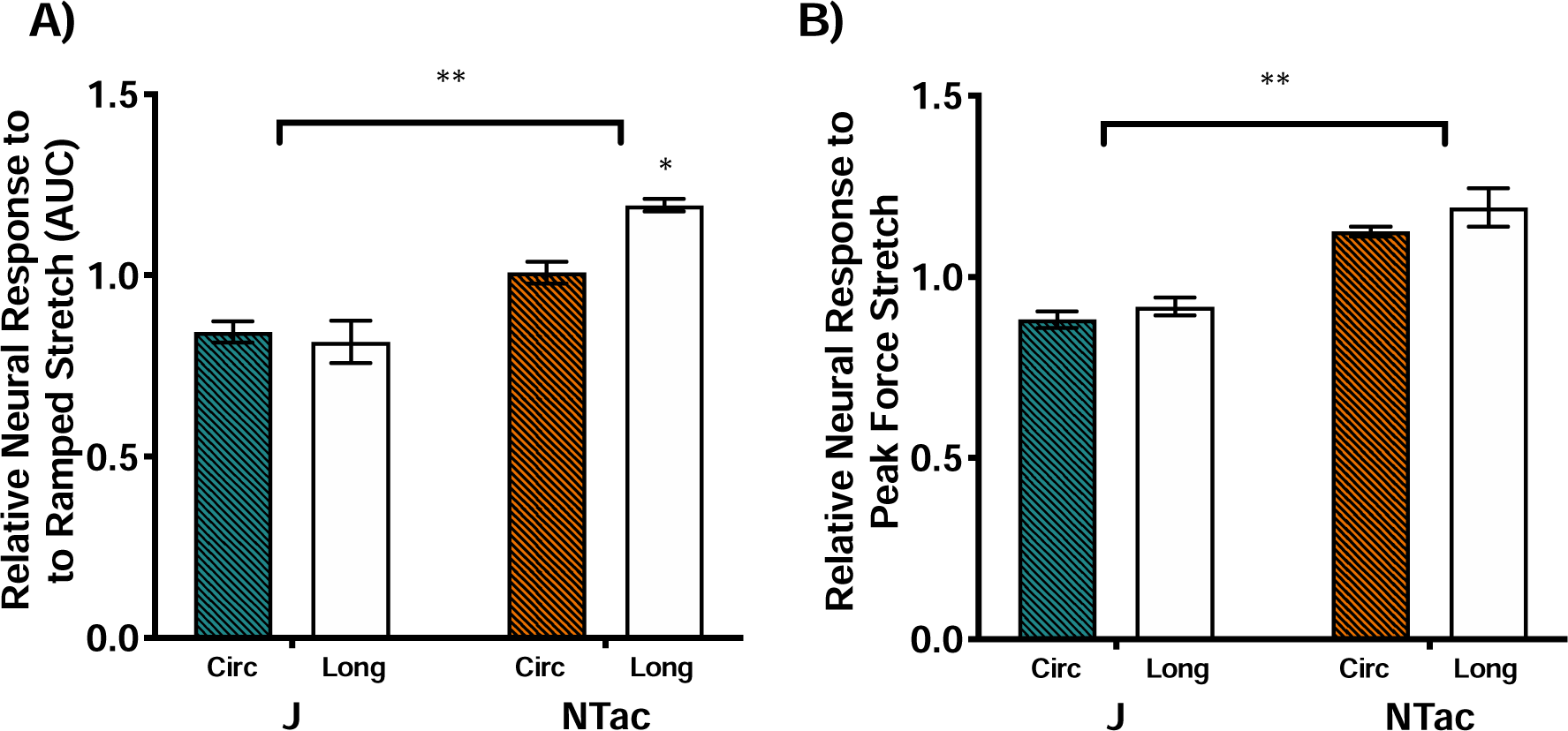
Colorectal spinal afferent responses to colorectal stretch measured by single-fiber recordings with an *ex vivo* colorectum-pelvic nerve preparation. In ZYM-treated BL/6NTac and BL/6J mice, the stimulus-response function (SRF) was measured before, after AVP application to the receptive field, and subsequent washout. **(A)** A 2 (Strain) x 2 (Stretch Direction) ANOVA revealed a significant main effect of Strain (F (1, 15) = 30.810, p < 0.001) and a Strain x Stretch Direction interaction (F (1, 15) = 4.787, p < 0.05) on AUC.). **(B)** Similarly, 2 x 2 ANOVA for response during peak stretch force application confirmed a significant main effect of Strain (1, 15) = 57.203, p < 0.001), confirming a greater response magnitude in BL/6NTac mice treated with ZYM compared to BL/6J mice treated with ZYM. No other significant main effects or interactions were observed (all p< 0.05).

### AVPR1A/*AVPR1A* expression in colorectal biopsies from clinical cohorts with and without persistent abdominal pain and healthy controls

In support of the experimental data described above, the clinical evaluation of *AVPR1A* expression was explored in two cohorts of patients with varying self-reported intensities of abdominal pain. In **Figure 5A**, pediatric patients (males and females, age 7-17) undergoing diagnostic colorectal biopsy for abdominal pain were compared to pediatric patients undergoing colonoscopy for functional nausea or painless non-cancerous polyp monitoring (n=3 patients). Patients with Rome III criteria for IBS (and controls) in the absence of other diagnoses (e.g., active *Helicobacter pylori* infection and/or IBD) or clinical signs of disease were assessed using the Pain Burden Inventory (PBI), in which patients self-reported their pain severity. Relative *AVPRA* mRNA expression (2^-ddCt^ method) comparing IBS patients with low and high self-reported pain [defined as a PBI score of 0-2 (low pain, n=13) or > 2 (high pain, n=25), respectively]. The patients reporting high pain burden appeared to have higher *AVPR1A* expression than IBS patients reporting low pain burden or healthy controls, though this difference was not significant (**Figure 5A**). The relatively high variability may be due to the relatively small number of patients in the control group (n = 3) and difficulty recruiting this population (as screening colonoscopy is not indicated for pediatric patients).

**Figure 5.**
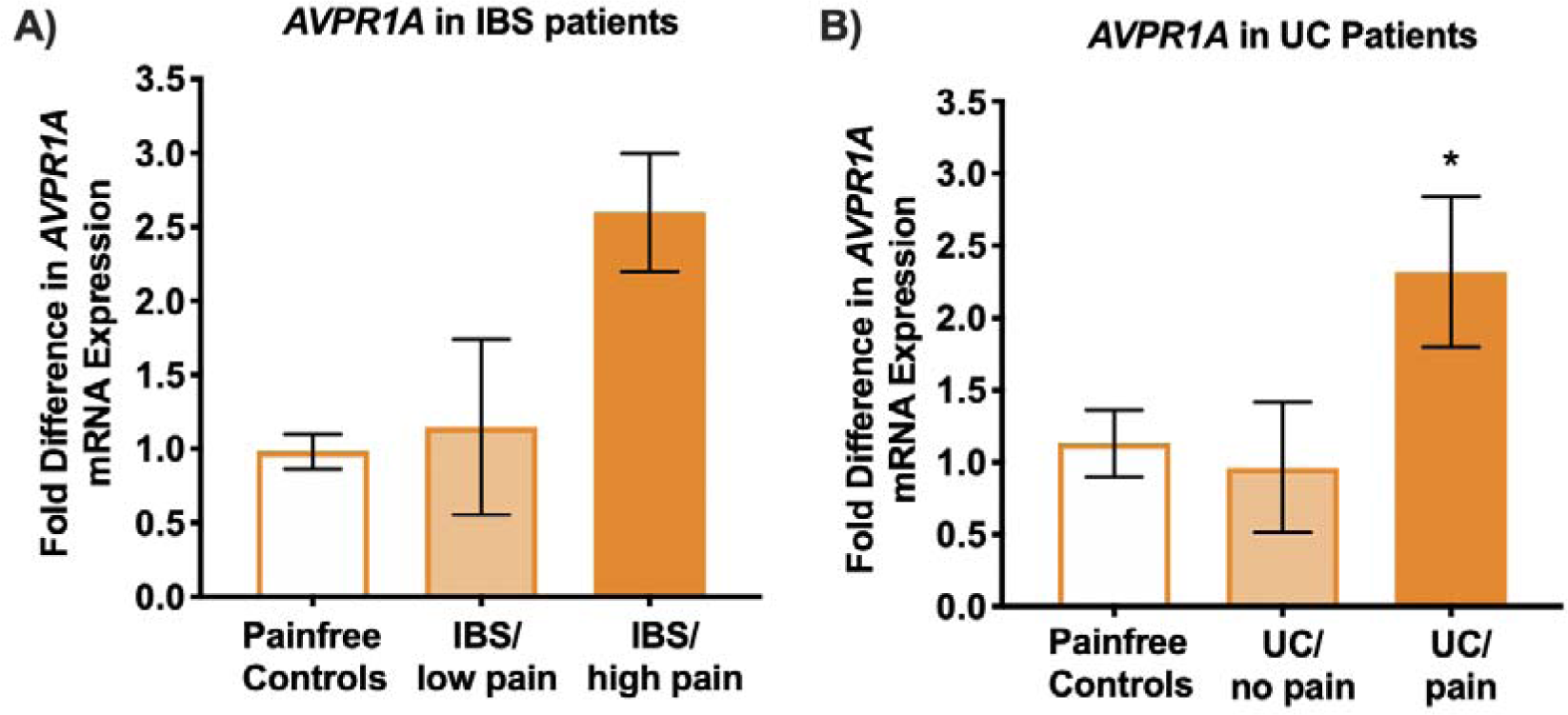
**(A)** *AVPR1A mRNA* levels from pediatric IBS patients’ colonic biopsies with low or high levels of reported pain compared to pain-free controls. IBS/FAP (n = 19) vs. Functional Nausea/painless rectal bleeding as controls (n = 3). T statistic = 1.462, p value = 0.159, MEANS: IBS/FAP=2.58 (SEM=0.334) vs control = 1.262 (SEM=0.545). This could be due to a low number of controls and difficulty finding appropriate controls given that preventative colonoscopy is not recommended until adulthood, and these are all adolescents. **(B)** *AVPR1A mRNA* levels from adult patients with active UC who report pain compared to those reporting no pain and pain-free controls. n = 6-9 per group. A 2 x 2 ANOVA showed a main effect of pain status but not disease status. Active/NP = 0.51 ((SEM = 0.162), Active/P = 2.23 (0.467), Inactive/P = 1.56 (0.339), Inactive/NP = 1.11 (0.302). * Indicate significant ANOVA, p < 0.05.

In a second study, *AVPR1A* expression was assessed in rectal biopsies from cohorts of adults with active (n = 6 with no pain, 9 with pain) or inactive (n = 6 with no pain, 7 with pain) ulcerative colitis (UC, as well as healthy pain-free controls (n = 6). Relative mRNA expression (2^-ddCt^ method) values were analyzed using 2 (Active vs. Inactive) x 2 (Pain vs No Pain) ANOVA which revealed a significant main effect of pain status (F (1,24) = 6.935, p = 0.015) but no other significant main effects or interactions (all p > 0.05). *AVPR1A* expression was significantly higher in biopsies from patients with pain than those without pain, regardless of their UC disease activity status (**Figure 5B**).

### Immunohistochemistry staining reveals possible co-localization between AVPR1A and neuronal terminal endings

Co-labeling of the distal colon with antisera against PGP9.5 (neuronal marker) and AVPR1A revealed overlap of the two markers, suggesting AVPR1A is expressed in neurons with terminals in the colon wall (**Figure 6**). PGP9.5 labeling does not discriminate between extrinsic primary afferents (DRG) neurons) and neurons of the enteric nervous system, but we have already shown that *Avpr1a/*AVPR1A expression is increased in the colorectum but not in colon-specific DRG neurons (data not shown) and ZYM- induced increases in *Avpr1a/*AVPR1A were restricted to colonic tissues (refer to **Figure 3**).

**Figure 6.**
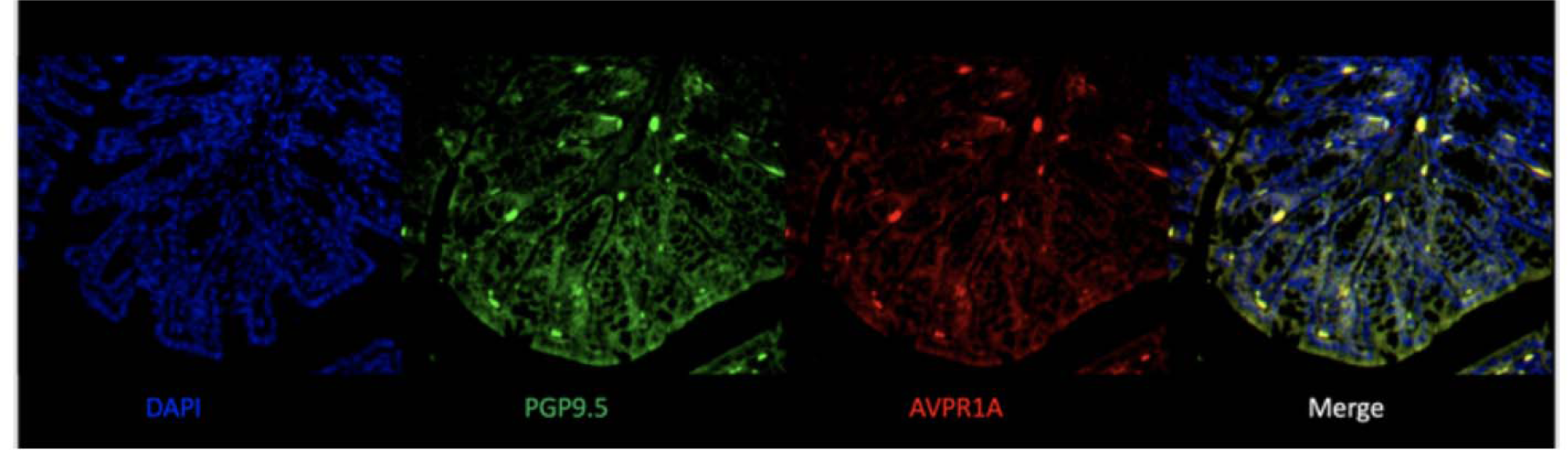
Representative images of immunohistochemistry stains of distal colon cross-sections from BL/6NTac mice instilled with ZYM. Sections were stained with DAPI nuclear stain (blue), AVPR1A (red), PGP9.5 for neurons (green), and merge of all three to show co-localization of AVPR1A and neuronal cell bodies (yellow), n=5 BL/6NTac-ZYM.

### *In vitro* Ca^2+^ imaging of ENS neurons from VH mice exhibited significantly greater Ca2+ influx to AVP

We measured Ca^2+^ transients in separate cultures of extrinsic colon specific DRG and enteric neurons in response to 1) the TRPV1 agonist capsaicin, and 2) the AVPR1A agonist, AVP. When comparing neurons from BL/6NTac mice with ZYM-VH to SAL-treated controls, neither DRG nor ND ganglion neurons retrogradely labeled from the colon exhibited altered cellular Ca^2+^ influx (ΔF) or changes in the % of cells responding to any chemical stimulus (**Supplemental Figure 8**) (all p > 0.05). Enteric neurons isolated from the myenteric plexus exhibited increased responses (ΔF) to AVP (p < 0.001) and capsaicin (p < 0.001) (**Figure 7**) suggesting enteric neuron hyperresponsiveness may play a role in increased sensory neuronal response to stretch (in line with single fiber teasing experiments) and VH in BL/6NTac treated with ZYM (**Figure 4**).

**Figure 7.**
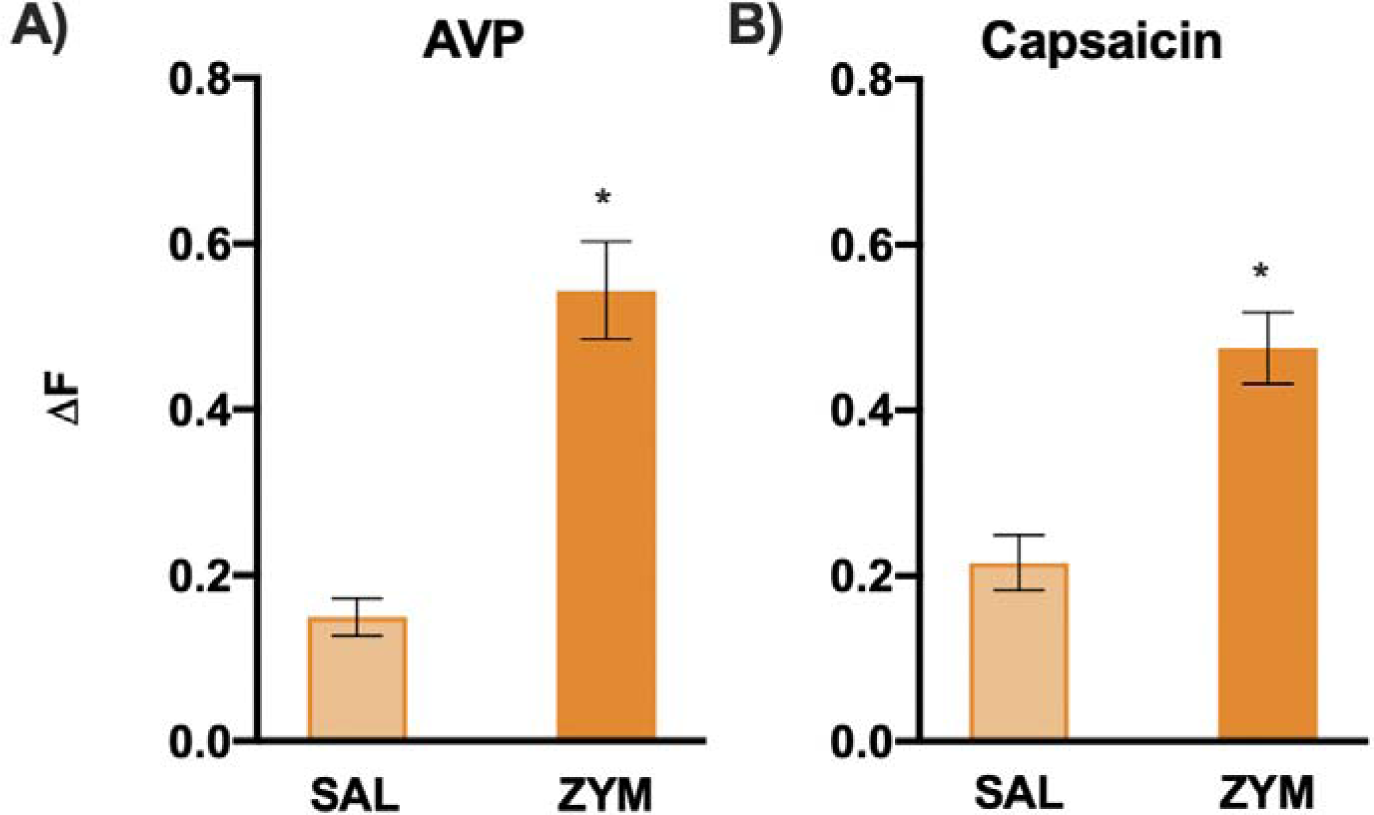
**A)** In vitro Ca^2+^ imaging of ENS neurons revealed individual neurons from VH mice (Mean = 0.5342, SEM = 0.0671) exhibited significantly greater Ca^2+^ influx (ΔF) in response to AVP compared to SAL controls (Mean = 0.2282, SEM = 0.0240). p<0.001. A) ENS neurons revealed individual neurons from VH mice (Mean = 0.2155, STEM= 0.0243) exhibited greater Ca^2+^ influx (DF) in response to capsaicin compared to SAL controls (Mean = 0.0228, SEM = 0.0212). The % of neurons responding to AVP (Person Chi-Square p = 0.051) and capsaicin (Pearson Chi-Square p = 0.569) were not statistically significant (data not shown).

## Discussion

The C57BL/6 inbred strain is ubiquitous in biomedical research serving as the primary reference or “normal” comparison strain for most mouse behavior and for the mouse genome assembly. However, the term BL/6 is often used to refer to a collection of highly genetically similar, but not identical, strains of mice [47], with little acknowledgement of the genotypic and phenotypic variations that exist between the closely related substrains. Inbred mouse strains are defined by the assumption of genome wide homozygosity for all individual members and by the assumption of genomic fixation across time. Each BL/6 substrain satisfies these expectations but multiple published studies comparing substrains have reported variability across various phenotypes, including motor coordination, behavior, and pain sensitivity[48–52]. Only recently has the differential pain susceptibility across BL/6 strains been explored[53–58]. Along these lines, we identified two C57BL/6 substrains with differential susceptibility to the development of VH and leveraged whole genome sequencing[23, 46] to identify *Avpr1a* as the highest priority candidate gene for differential VH susceptibility. AVPR1A belongs to a subfamily of G-protein coupled receptors (GPCRs), and even though GPCRs have been found to modulate neuronal sensitization of the gut, the role of AVPR1A and its agonists have not previously been explored in basal colonic function or in VH susceptibility.

In line with prior work on the role of *Avpr1a* in somatic pain[59], the SNP differentiating BL/6J and BL/6NTac mice is in a proposed 5’ regulatory region outside of the *Avpr1a* coding sequence, pointing to a potential effect on gene expression rather than alterations in the encoded protein product. We confirmed a strain-dependent mRNA expression difference corresponding to VH susceptibility and a corresponding strain difference in functional sensitivity in response to application of exogenous arginine-vasopressin (AVP), the AVPR1A agonist, via single-fiber recordings in an *ex vivo* colon-pelvic nerve single fiber recording preparation. Moreover, *AVPR1A* expression was also higher in colorectal biopsies collected from UC patients with persistent abdominal pain when compared to disease-severity matched UC patients with no pain, supporting the translational potential of AVPR1A genotype/expression as a risk factor for development of VH.

While *Avpr1a*/*AVPR1A* has been associated with somatic pain variability in prior work, the relationship between *Avpr1a* expression and somatic inflammatory pain was opposite to what we report for visceral pain. We found that increased *Avpr1a* expression is associated with *hypo*sensitivity for somatic pain, an effect that is sex- and stress-dependent in animals and human subjects [44, 59, 60]. While these studies did not assess *Avpr1a* expression in all tissues comprising of the pain processing pathway, from area of inflammation to the brain, *Avpr1a* expression in the mouse DRG was inversely correlated with pain behavior (i.e., lower *Avpr1a* expression in pain sensitive mice). In contrast, the current data indicate that differential pain sensitivity between strains is not correlated with *Avpr1a* expression in DRG. It has also been previously shown that *Avpr1a* has some expression in small diameter DRG neurons (referred to as putative nociceptors in the report) and that expression was increased in naïve mice with SNPs corresponding to higher expression of *Avpr1a* and lower pain intensity[59]. Conversely, our data indicate that *Avpr1a* expression in enteric neurons of the colon is higher in VH- susceptible mice and that these neurons are more responsive to stimulation with the agonist AVP. The disagreement in the direction and location of prior findings and ours is likely due to the unique functional aspects of the distal colon compared to skin/distal somatic tissues. Relevant to our findings, *Avpr1a* may represent a novel point of convergence between VH following transient gastroenteritis and stress-induced VH that could explain the stress-induced pain exacerbations that many DGBI patients experience. Along these lines, endogenous vasopressin enters circulation downstream of the stress response and subsequent AVPR1A activation in the bowel results in vasoconstriction, alterations in water absorption in the colon, and abdominal pain/fecal urgency[61–63], mimicking the effects of exogenous vasopressin administration in the clinical setting (e.g., to control GI bleeding, etc.). For individuals with elevated colonic *Avpr1a/AVPR1A* expression and VH, vasopressin released in response to stress could result in significant exacerbation of VH through binding of AVP to AVPR1A, thereby linking this receptor to a potential mechanism for the clinically recognized stress-induced exacerbations of abdominal pain seen in IBS and other DGBIs.

Even though previous work suggests neuronal plasticity and broad alteration to the gut-brain axis has been emphasized as an underlying factor in chronic abdominal pain, much of the previous literature has focused on DRG originating in pelvic, splanchnic, and ND ganglia due to the well-known relationship between visceral DRG sensitization and hypersensitivity [16, 64, 65]. More recently, however, published data suggest a possible role for the ENS in pathological pain[65–67], though the mechanism of communication between the ENS and DRGs remains incompletely understood. Based on the observations that neither spinal nor ND DRGs were hyper-responsive to chemical stimulation of capsaicin and AVP in the absence of enteric neurons but did exhibit increased stretch-responsiveness when functionally connected to the ENS in the *ex vivo* fiber teasing preparation, we propose that communication between the ENS and DRG may play a critical role in the development of VH. Understanding a precise mechanism for enteric cell involvement in abdominal pain needs further evaluation. However, these data point to the potential utility of targeting *Avpr1a*/AVPR1A expression and/or function locally in the colon in the prevention and/or treatment of VH We propose *Avpr1a* as a novel colon-specific therapeutic target for VH based on our rigorously collected preliminary data incorporating genomic comparisons, transcript covariance analysis, and functional sensitivity to pharmacological manipulation. *Avpr1a* represents a potential biomarker for VH susceptibility with implications for the etiology of IBS and other DGBIs conditions as well as for novel therapeutic interventions targeting this specific underlying mechanism.

The present study has certain limitations that should be noted. One notable constraint is that the experimental investigations were conducted solely using male mouse models, thereby limiting our ability to explore potential sex-specific differences. Consequently, caution must be exercised when extrapolating the study’s conclusions to female populations, as biological variations between male and female mice may exist and could impact the results. In addition, while there is no ongoing inflammation, the ZYM model may be considered a model of post-inflammatory VH and whether this exact mechanism plays a role in individual differences in susceptibility to other forms of VH is yet to be determined. Furthermore, our immunobiological staining does not discriminate the location of AVPR1A in the distal colon between DRG endings and the enteric neurons, so our future work will include higher-resolution exploration of AVPR1A location. Lastly, we would like to acknowledge there needs to be more literature of human diseases or traits associated with this gene (in general, not just pain-related), so the unavailability of a replication, combined with male-only association is another potential limitation here. Therefore, with all the publicly available data now, a thoughtfully designed analysis should be able to address this potential limitation later down the road.

## Acknowledgments

We want to thank our primary funding sources for this work: A recruitment enhancement funds (EEY and KMB) and a Developmental Research Project Program (DRPP, EEY) award through the Kansas-INBRE Institutional Development Award (IDeA) from the National Institute of General Medical Sciences of the National Institutes of Health [P20 GM103418]. We want to thank Dr. Gerald Gebhart for his contributions to the initial studies related to our preclinical model and for offering feedback on the drafted manuscript. We also thank Dr. Milad Mortazavi and Dr. Abraham Palmer for collaborating and publishing whole genome sequencing data between BL/6 and BL/10 mouse substrain models.

## References

1. Mayer, E.A. and G.F. Gebhart, Basic and clinical aspects of visceral hyperalgesia. Gastroenterology, 1994. 107(1): p. 271–93.

2. Camilleri, M., S.B. Saslow, and A.E. Bharucha, Gastrointestinal sensation. Mechanisms and relation to functional gastrointestinal disorders. Gastroenterol Clin North Am, 1996. 25(1): p. 247–58.

3. Camilleri, M., B. Coulie, and J.F. Tack, Visceral hypersensitivity: facts, speculations, and challenges. Gut, 2001. 48(1): p. 125–31.

4. Barbara, G., et al., New pathophysiological mechanisms in irritable bowel syndrome. Aliment Pharmacol Ther, 2004. 20 **Suppl 2**: p. 1–9.

5. Sayuk, G.S., et al., Opioid medication use in patients with gastrointestinal diagnoses vs unexplained gastrointestinal symptoms in the US Veterans Health Administration. Aliment Pharmacol Ther, 2018. 47(6): p. 784–791.

6. Hser, Y.I., et al., Chronic pain among patients with opioid use disorder: Results from electronic health records data. J Subst Abuse Treat, 2017. 77: p. 26–30.

7. Zhou, Q. and G.N. Verne, New insights into visceral hypersensitivity--clinical implications in IBS. Nat Rev Gastroenterol Hepatol, 2011. 8(6): p. 349–55.

8. Azpiroz, F., et al., Mechanisms of hypersensitivity in IBS and functional disorders. Neurogastroenterol Motil, 2007. 19(1 Suppl): p. 62-88.

9. Clouse, R.E., et al., Functional abdominal pain syndrome. Gastroenterology, 2006. 130(5): p. 1492–7.

10. Morris, G.P., et al., Hapten-induced model of chronic inflammation and ulceration in the rat colon. Gastroenterology, 1989. 96(3): p. 795–803.

11. Sengupta, J.N., et al., Effects of kappa opioids in the inflamed rat colon. Pain, 1999. 79(2-3): p. 175–85.

12. Lu, G., et al., Inflammation modulates in vitro colonic myoelectric and contractile activity and interstitial cells of Cajal. Am J Physiol, 1997. 273(6): p. G1233–45.

13. Verma-Gandhu, M., et al., Visceral pain perception is determined by the duration of colitis and associated neuropeptide expression in the mouse. Gut, 2007. 56(3): p. 358–64.

14. Zhang, H.Q., et al., Therapeutic effects of Clostridium butyricum on experimental colitis induced by oxazolone in rats. World J Gastroenterol, 2009. 15(15): p. 1821–8.

15. Burton, M.B. and G.F. Gebhart, Effects of intracolonic acetic acid on responses to colorectal distension in the rat. Brain Res, 1995. 672(1-2): p. 77–82.

16. Kamp, E.H., et al., Quantitative assessment and characterization of visceral nociception and hyperalgesia in mice. Am J Physiol Gastrointest Liver Physiol, 2003. 284(3): p. G434–44.

17. Coutinho, S.V., S.T. Meller, and G.F. Gebhart, Intracolonic zymosan produces visceral hyperalgesia in the rat that is mediated by spinal NMDA and non-NMDA receptors. Brain Res, 1996. 736(1-2): p. 7–15.

18. Liu, S.B., et al., Long-term upregulation of cortical glutamatergic AMPA receptors in a mouse model of chronic visceral pain. Mol Brain, 2015. 8(1): p. 76.

19. Bryant, C.D., The blessings and curses of C57BL/6 substrains in mouse genetic studies. Ann N Y Acad Sci, 2011. 1245: p. 31–3.

20. Young, E.E., et al., Systems genetic and pharmacological analysis identifies candidate genes underlying mechanosensation in the von Frey test. Genes Brain Behav, 2016. 15(6): p. 604–615.

21. Zurita, E., et al., Genetic polymorphisms among C57BL/6 mouse inbred strains. Transgenic Res, 2011. 20(3): p. 481–9.

22. Churchill, G.A., et al., The Collaborative Cross, a community resource for the genetic analysis of complex traits. Nat Genet, 2004. 36(11): p. 1133–7.

23. Mortazavi, M., et al., SNPs, short tandem repeats, and structural variants are responsible for differential gene expression across C57BL/6 and C57BL/10 substrains. Cell Genom, 2022. 2(3).

24. Lacy, B.E. and N.K. Patel, Rome Criteria and a Diagnostic Approach to Irritable Bowel Syndrome. J Clin Med, 2017. 6(11).

25. Meller, S.T. and G.F. Gebhart, Intraplantar zymosan as a reliable, quantifiable model of thermal and mechanical hyperalgesia in the rat. Eur J Pain, 1997. 1(1): p. 43–52.

26. Christianson, J.A. and G.F. Gebhart, Assessment of colon sensitivity by luminal distension in mice. Nat Protoc, 2007. 2(10): p. 2624–31.

27. Feng, B., P.R. Brumovsky, and G.F. Gebhart, Differential roles of stretch-sensitive pelvic nerve afferents innervating mouse distal colon and rectum. Am J Physiol Gastrointest Liver Physiol, 2010. 298(3): p. G402–9.

28. Schneider, C.A., W.S. Rasband, and K.W. Eliceiri, NIH Image to ImageJ: 25 years of image analysis. Nat Methods, 2012. 9(7): p. 671–5.

29. Vilz, T.O., et al., Functional assessment of intestinal motility and gut wall inflammation in rodents: analyses in a standardized model of intestinal manipulation. J Vis Exp, 2012(67).

30. Schwartz, E.S., et al., Synergistic role of TRPV1 and TRPA1 in pancreatic pain and inflammation. Gastroenterology, 2011. 140(4): p. 1283–1291 e1-2.

31. Bolon, B., I.D. Pardo, and G.J. Krinke, The Science and Art of Nerve Fiber Teasing for Myelinated Nerves: Methodology and Interpretation. Toxicol Pathol, 2020. 48(1): p. 49–58.

32. Deberry, J.J., et al., Abdominal pain and the neurotrophic system in ulcerative colitis. Inflamm Bowel Dis, 2014. 20(12): p. 2330–9.

33. Grossi, V., et al., Characterizing Clinical Features and Creating a Gene Expression Profile Associated With Pain Burden in Children With Inflammatory Bowel Disease. Inflamm Bowel Dis, 2020. 26(8): p. 1283–1290.

34. Malin, S.A., et al., TPRV1 expression defines functionally distinct pelvic colon afferents. J Neurosci, 2009. 29(3): p. 743–52.

35. Zhang, Y. and W. Hu, Mouse enteric neuronal cell culture. Methods Mol Biol, 2013. 1078: p. 55–63.

36. Zhang, Y. and W. Hu, Mouse Enteric Neuronal Cell Culture. Methods Mol Biol, 2021. 2311: p. 63–71.

37. Pierce, A.N., et al., Neonatal vaginal irritation results in long-term visceral and somatic hypersensitivity and increased hypothalamic-pituitary-adrenal axis output in female mice. Pain, 2015. 156(10): p. 2021–2031.

38. Feng, B., et al., Long-term sensitization of mechanosensitive and -insensitive afferents in mice with persistent colorectal hypersensitivity. Am J Physiol Gastrointest Liver Physiol, 2012. 302(7): p. G676–83.

39. Greco, V., et al., Histochemistry of the colonic epithelial mucins in normal subjects and in patients with ulcerative colitis. A qualitative and histophotometric investigation. Gut, 1967. 8(5): p. 491–6.

40. Simon, M.M., et al., A comparative phenotypic and genomic analysis of C57BL/6J and C57BL/6N mouse strains. Genome Biol, 2013. 14(7): p. R82.

41. Warden, A.S., et al., Inbred Substrain Differences Influence Neuroimmune Response and Drinking Behavior. Alcohol Clin Exp Res, 2020. 44(9): p. 1760–1768.

42. Mekada, K., et al., Genetic differences among C57BL/6 substrains. Exp Anim, 2009. 58(2): p. 141–9.

43. Wilson, S.G., et al., Identification of quantitative trait loci for chemical/inflammatory nociception in mice. Pain, 2002. 96(3): p. 385–391.

44. Nair, H.K., et al., Genomic loci and candidate genes underlying inflammatory nociception. Pain, 2011. 152(3): p. 599–606.

45. Mogil, J.S., et al., Pain sensitivity and vasopressin analgesia are mediated by a gene-sex-environment interaction. Nat Neurosci, 2011. 14(12): p. 1569–73.

46. Mortazavi, M., et al., Polymorphic SNPs, short tandem repeats and structural variants are responsible for differential gene expression across C57BL/6 and C57BL/10 substrains. bioRxiv, 2021: p. 2020.03.16.993683.

47. Mouse Genome Sequencing, C., et al., Initial sequencing and comparative analysis of the mouse genome. Nature, 2002. 420(6915): p. 520-62.

48. Hovland, D.N., Jr., et al., Identification of a murine locus conveying susceptibility to cadmium-induced forelimb malformations. Genomics, 2000. 63(2): p. 193–201.

49. Egan, C.M., et al., Recurrent DNA copy number variation in the laboratory mouse. Nat Genet, 2007. 39(11): p. 1384–9.

50. Aguirre, J.E., J.H. Winston, and S.K. Sarna, Neonatal immune challenge followed by adult immune challenge induces epigenetic-susceptibility to aggravated visceral hypersensitivity. Neurogastroenterol Motil, 2017. 29(9).

51. Tran, L., et al., Importance of epigenetic mechanisms in visceral pain induced by chronic water avoidance stress. Psychoneuroendocrinology, 2013. 38(6): p. 898–906.

52. Mahurkar-Joshi, S. and L. Chang, Epigenetic Mechanisms in Irritable Bowel Syndrome. Front Psychiatry, 2020. 11: p. 805.

53. Ulker, E., et al., C57BL/6 substrain differences in formalin-induced pain-like behavioral responses. Behav Brain Res, 2020. 390: p. 112698.

54. Bryant, C.D., et al., C57BL/6 substrain differences in inflammatory and neuropathic nociception and genetic mapping of a major quantitative trait locus underlying acute thermal nociception. Mol Pain, 2019. 15: p. 1744806918825046.

55. Leo, S., et al., *Differences in nociceptive behavioral performance between C57BL/6J*, *129S6/SvEv, B6 129 F1 and NMRI mice*. Behav Brain Res, 2008. 190(2): p. 233-42.

56. Mekada, K. and A. Yoshiki, Substrains matter in phenotyping of C57BL/6 mice. Exp Anim, 2021.

57. Bryant, C.D., et al., Behavioral differences among C57BL/6 substrains: implications for transgenic and knockout studies. J Neurogenet, 2008. 22(4): p. 315–31.

58. Smith, J.C., A Review of Strain and Sex Differences in Response to Pain and Analgesia in Mice. Comp Med, 2019. 69(6): p. 490–500.

59. Mogil, J.S., et al., Pain sensitivity and vasopressin analgesia are mediated by a gene-sex-environment interaction. Nat Neuroscie, 2011. 14(12): p. 1569–1573.

60. Roach, K.L., et al., The AVPR1A Gene and Its Single Nucleotide Polymorphism rs10877969: A Literature Review of Associations with Health Conditions and Pain. Pain Manag Nurs, 2018. 19(4): p. 430–444.

61. Xu, X., D.L. Brining, and J.D. Chen, Effects of vasopressin and long pulse-low frequency gastric electrical stimulation on gastric emptying, gastric and intestinal myoelectrical activity and symptoms in dogs. Neurogastroenterol Motil, 2005. 17(2): p. 236–44.

62. Caras, S.D., et al., The effect of intravenous vasopressin on gastric myoelectrical activity in human subjects. Neurogastroenterol Motil, 1997. 9(3): p. 151–6.

63. Conn, H.O., et al., Intraarterial vasopressin in the treatment of upper gastrointestinal hemorrhage: a prospective, controlled clinical trial. Gastroenterology, 1975. 68(2): p. 211–21.

64. Feng, B. and G.F. Gebhart, Characterization of silent afferents in the pelvic and splanchnic innervations of the mouse colorectum. Am J Physiol Gastrointest Liver Physiol, 2011. 300(1): p. G170–80.

65. Brierley, S.M., et al., Splanchnic and pelvic mechanosensory afferents signal different qualities of colonic stimuli in mice. Gastroenterology, 2004. 127(1): p. 166–78.

66. Holland, A.M., et al., The enteric nervous system in gastrointestinal disease etiology. Cell Mol Life Sci, 2021. 78(10): p. 4713–4733.

67. Lucarini, E., et al., Role of Enteric Glia as Bridging Element between Gut Inflammation and Visceral Pain Consolidation during Acute Colitis in Rats. Biomedicines, 2021. 9(11).

