## supplemental Figures for "Identification of arginine-vasopressin receptor 1a (Avpr1a/AVPR1A) as a novel candidate gene for chronic visceral pain"

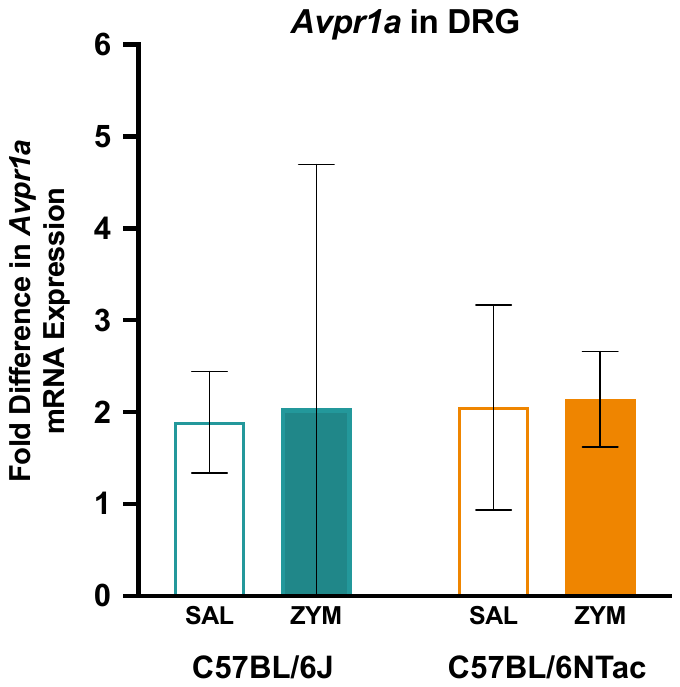

**Supplemental Figure 1.** Dorsal root ganglion (DRG) *Avpr1a* mRNA levels were measured in SAL or ZYM-treated BL/6NTac and BL/6J mice (all p > 0.05).

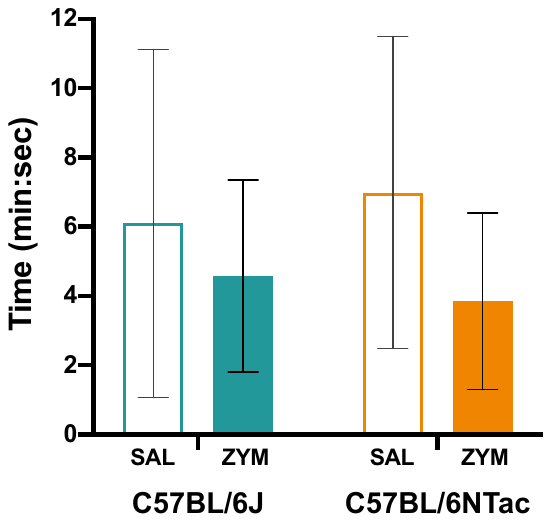

**Supplemental Figure 2**. **Colonic transit (CT) assay reveals no differences in motility time regardless of strain (BL/6NTac vs. BL/6J) or condition (SAL vs. ZYM).** *Teal bars = J, Orange bars = N/Tac. Statistical analysis was measured using a 2x2 ANOVA. n=5/group. All p > 0.5 (p = 0.664mouse group; p=2.605 treatment; p =0.480 mouse group*treatment).

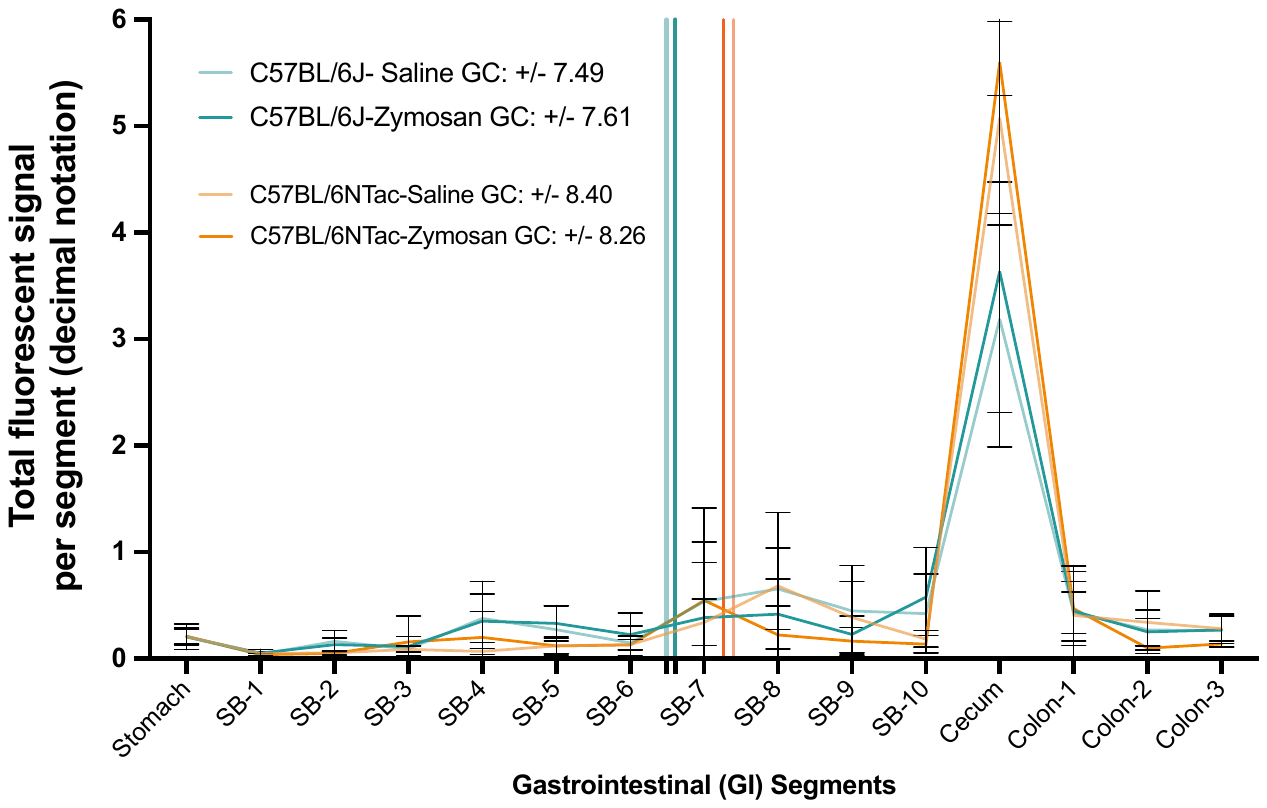

**Supplemental Figure 3**. **Global (GI) transit assay reveals no differences in motility measurement regardless of strain (BL/6J vs. BL/6NTac) or condition (Naïve/SAL vs. ZYM).** Statistical analysis was measured using a 2x2 ANOVA. n=5/group. All p > 0.5 (p = 0.977 treatment; p = 0.125 mouse group; p = 0.800 treatment*mouse group).

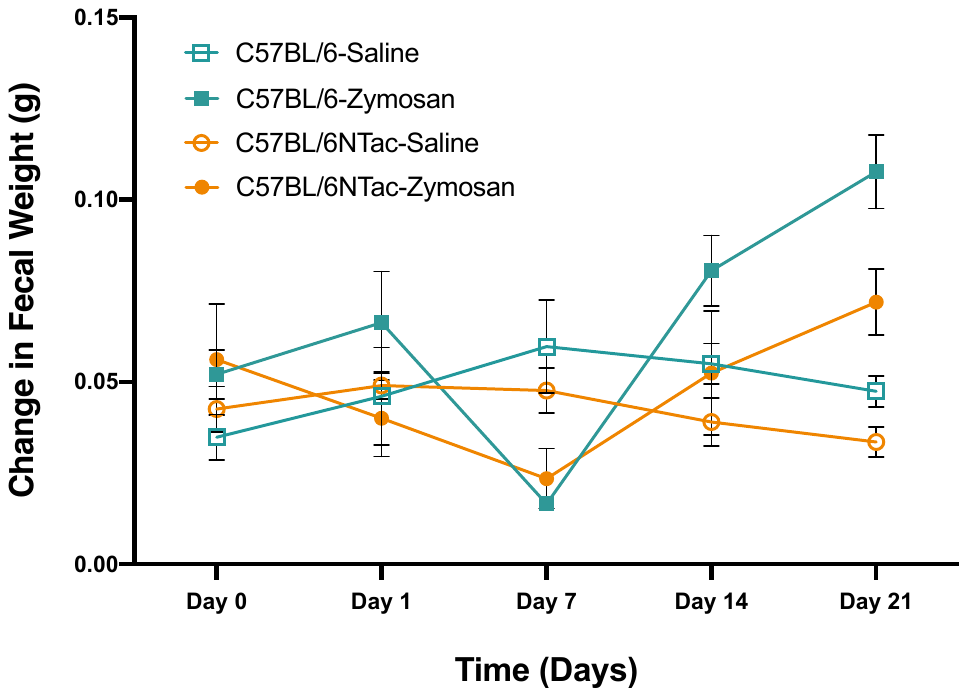

**Supplemental Figure 4**. Change in fecal water weight was measured by collecting fecal weight (4 pellets collected/mouse) at two time points: (1) immediately after a mouse excreted pellets (wet weight) and (2) after fecal matter had dried (dry weight) throughout VH development. Average weight differences (wet fecal matter – dry fecal matter) for each strain and condition are shown above. n=5/group. * Open Square Teal = BL/6J-SAL, Square Teal = BL/6J-ZYM, Open Orange Circle = BL/6NTac-SAL, Orange Circle = BL/6NTac-ZYM. Statistical analysis was measured using a 2x2 ANOVA.

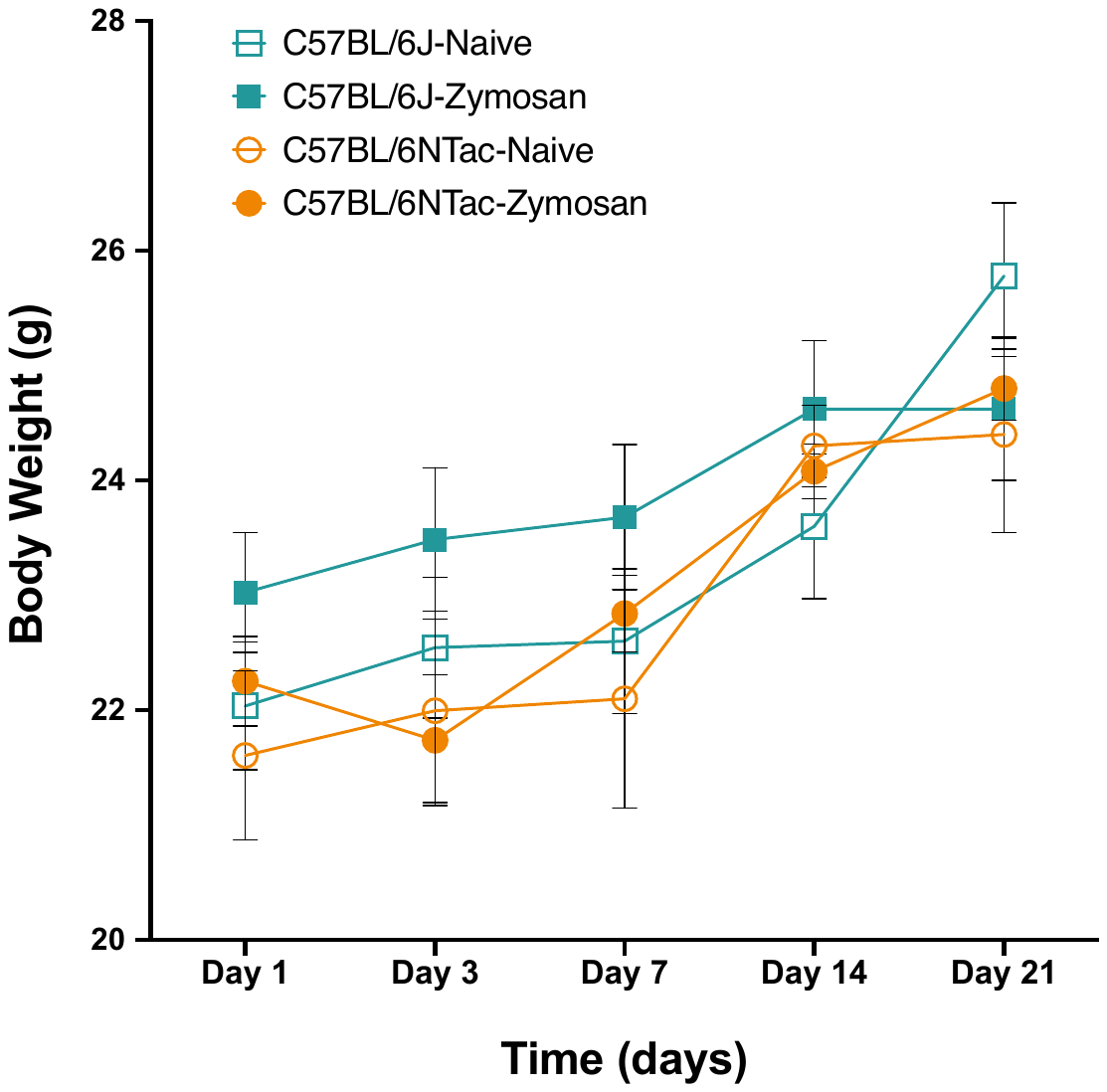

**Supplemental Figure 5**. **Average change in body weight after zymosan or saline installations during VH development is not different between each strain and condition. n=5/group.** *Teal = BL/6J, *Orange = BL6/NTac. n=5/group. Statistical analysis was measured using a 2x2 ANOVA. (p= 0.295 strain. p= 0.453 condition. P = 0.777 strain*condition).

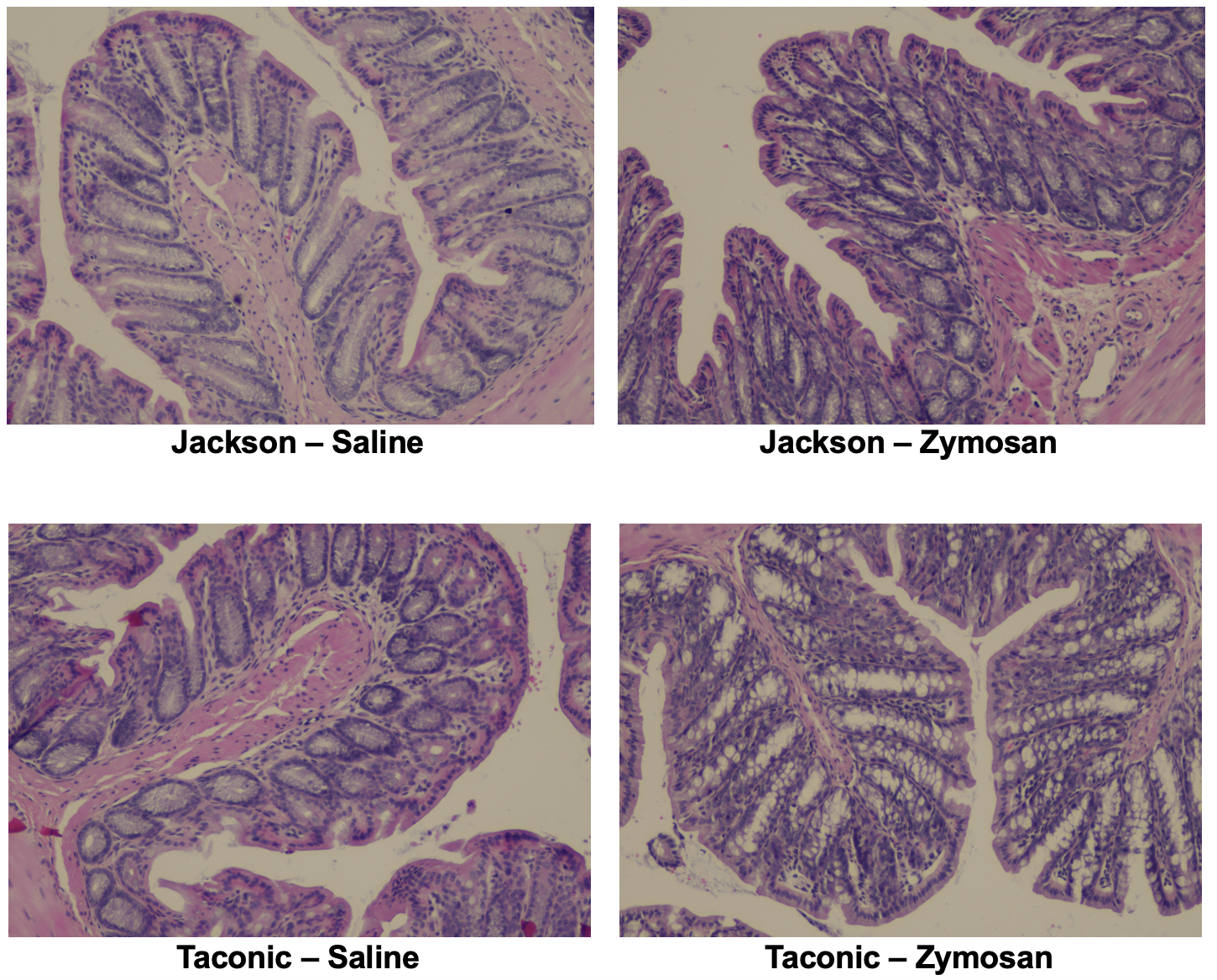

**Supplemental Figure 6**. Representative hematoxylin and eosin staining of the distal colon between strain (BL6/J vs. BL6/NTac) and condition (ZYM vs. SAL). n=5/group. No morphological and inflammatory nodules were present.

**A)	B)**

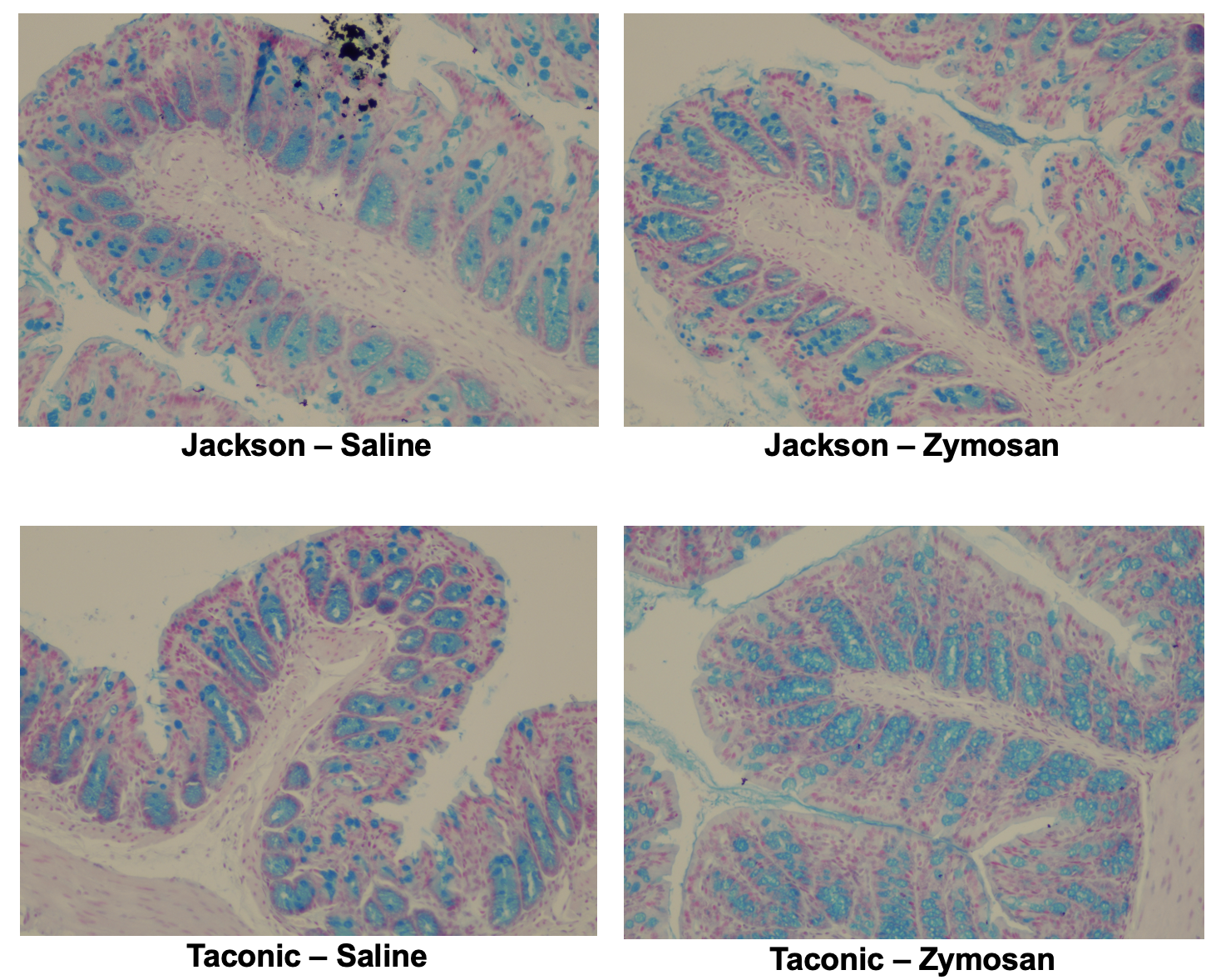

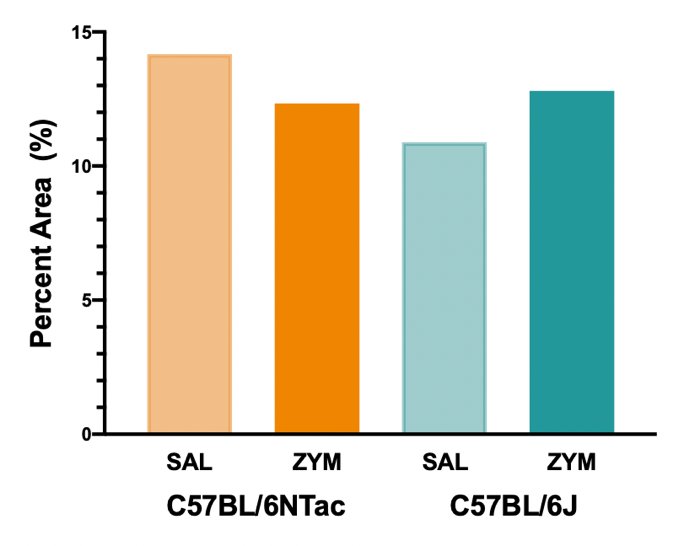

**Supplemental Figure 7. (A)** Representative Alcian blue stains for each strain (BL/6NTac vs. BL/J) and condition (ZYM vs. SAL). **(B)** Percent area of mucin cells is not significant between either strain or condition (ImageJ/FIJI). n=5/group. (p = 0.332 mouse group. p = 0.979 treatment. p = 0.201 mouse group*treatment).

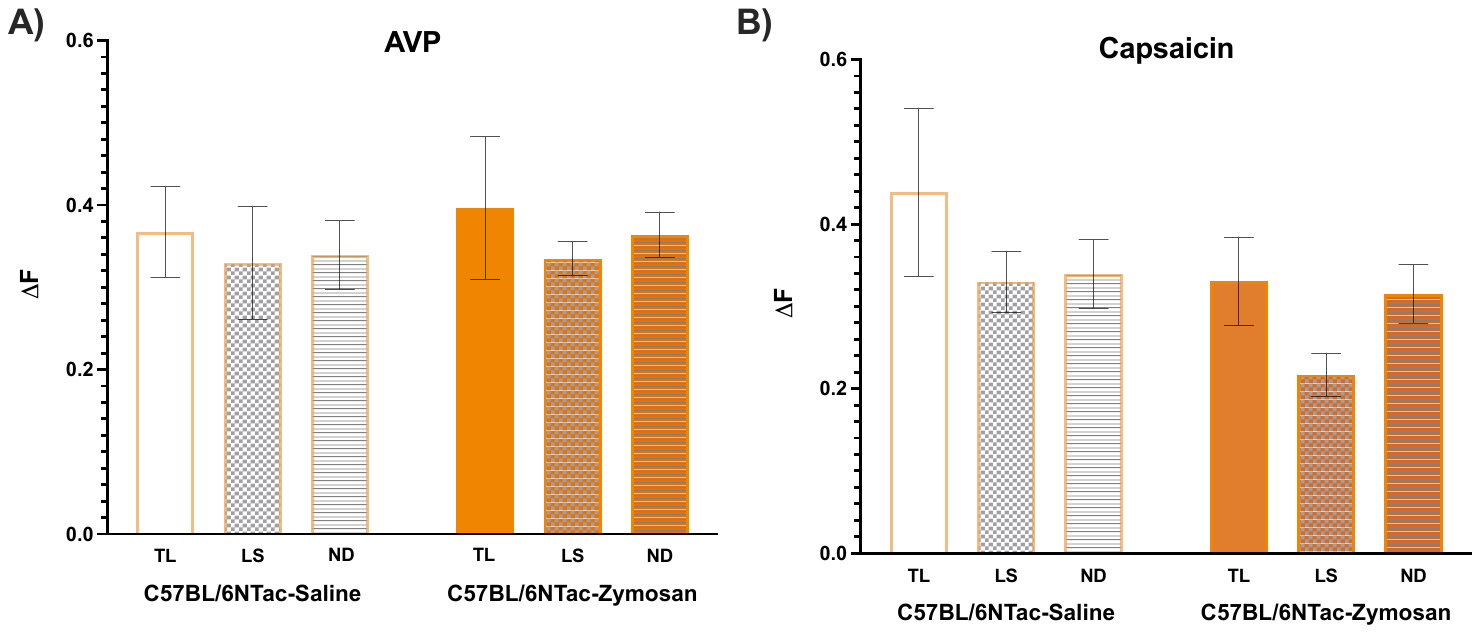

**Supplemental Figure 8**. In vitro Ca^2+^ imaging of retrogradely labeled extrinsic neurons revealed individual neurons from VH mice did not statistically differ in Ca^2+^ influx (ΔF) in response to AVP compared to SAL controls. n=5/group. (Capsaicin; p = 0.237. AVP; p = 0.486).
